# Pre-Injury Mechanoreceptor Ablation Reduces Nociceptor-Driven Spinal Cord Injury-Induced Neuropathic Pain

**DOI:** 10.1101/2022.12.18.520950

**Authors:** Christopher Sliwinski, Laura Heutehaus, Francisco J. Taberner, Lisa Weiss, Vasileios Kampanis, Bahardokht Tolou-Dabbaghian, Xing Cheng, Melanie Motsch, Paul A. Heppenstall, Rohini Kuner, Steffen Franz, Stefan G. Lechner, Norbert Weidner, Radhika Puttagunta

**Affiliations:** Laboratory of Experimental Neuroregeneration, Spinal Cord Injury Center, Heidelberg University Hospital, Heidelberg, Germany; Spinal Cord Injury Center, Heidelberg University Hospital, Heidelberg, Germany; Institute of Pharmacology, Heidelberg University, Im Neuenheimer Feld 366, 69120 Heidelberg, Germany; Department of Anesthesiology, University Medical Center Hamburg-Eppendorf, Martinistrasse 52, 20246 Hamburg, Germany; SISSA: Scuola Internazionale Superiore di Studi Avanzati, Via Bonomea, 265, 34136 Trieste, Italy

**Keywords:** spinal cord injury, dorsal root ganglion, dorsal horn, below level, neuropathic pain, TrkB, Aβ low-threshold mechanoreceptors, Aδ and C-fiber, peptidergic, nociceptors, mechanical allodynia, hypersensitivity, hyperexcitability

## Abstract

Evidence from previous studies supports the concept that spinal cord injury (SCI) induced neuropathic pain (NP) has its neural roots in the peripheral nervous system. There is uncertainty about how and to which degree nociceptors and mechanoreceptors contribute. Sensorimotor activation-based interventions (e.g. treadmill training) have been shown to reduce NP following experimental SCI, suggesting transmission of pain-alleviating signals through mechanoreceptors. At the same time, nociceptors have been shown to become hyperexcitable early after SCI and peptidergic axons sprout into deeper laminae of the below injury level dorsal horn. The aim of the present study is to comprehensively understand the relative contribution of each pathway in respect to NP presentation in a moderate mouse contusion SCI model. After genetic ablation of tropomyosin receptor kinase B (TrkB) expressing mechanoreceptors <u>before SCI</u> mechanical allodynia was reduced. The identical genetic ablation <u>after SCI</u> did not yield any change in pain behavior. CGRP sprouting into lamina III/IV below injury level as a consequence of SCI was not altered by either mechanoreceptor ablation. Moreover, detection of hyperexcitability in nociceptors, not in mechanoreceptors, in skin-nerve preparations of contusion SCI mice 7 days after injury makes a substantial direct contribution of mechanoreceptors to NP maintenance unlikely. SNS reporter mice allowing specific visualization of the entire nociceptor population confirmed significant sprouting of respective neurons into laminae III/IV as early as 5 days post-injury. Genetic ablation of SNS-Cre mice severely affected their overall health condition, which precluded them to undergo experimental SCI and subsequent further analysis. Complementing animal data, quantitative sensory testing in human SCI subjects indicated reduced mechanical pain thresholds, whereas the mechanical detection threshold was not altered. Taken together, early mechanoreceptor ablation modulates pain behavior, most likely through indirect mechanisms. Hyperexcitable nociceptors with consecutive peptidergic fiber sprouting in the dorsal horn are confirmed as the likely main driver of SCI-induced NP. Future studies need to focus on injury-derived factors triggering early onset nociceptor hyperexcitability, which could serve as targets for more effective therapeutic interventions.

## INTRODUCTION

Neuropathic pain (NP) represents a frequent concomitant condition in individuals suffering from spinal cord injury (SCI). Available treatments are insufficiently effective and frequently associated with significant side effects. Therefore, the identification of mechanisms underlying SCI-associated NP and subsequent development of targeted pharmacological and non-pharmacological interventions represent important aims to ultimately help improve the quality of life of SCI survivors.

Over many years of research, several neuroanatomical regions have been identified as potential sites of NP initiation and transmission. There is the actual injury site characterized by a variable degree of both overall spinal cord tissue damage and in particular a varying lesion with respect to ascending sensory pathways – the spinothalamic and the lemniscal tract. However, in recent years secondary alterations in the peripheral nervous system (PNS) below injury level have come more and more into focus. One key observation, confirmed in a number of studies, was the detection of nociceptor spontaneous activity and hyperexcitability found in dorsal root ganglia (DRG)-derived nociceptor cultures as well as analyses in skin-nerve preparations following experimental contusion SCI (1).

We wondered whether hyperexcitable nociceptors are no longer properly controlled by appropriate mechanoreceptor input due to SCI-induced para- or tetraparesis with resulting inability to stand or walk, thus depleting physiological sensory input from the skin and joints into the dorsal horn. Accordingly, a number of studies were conducted, which showed that sensorimotor activation through treadmill training in mice with moderate thoracic contusion SCI reduced pain behavior, which was associated with reversal of structural rearrangements below injury level in the dorsal horn, namely reduction of peptidergic nociceptor sprouting into laminae III-IV (2–4). These findings raised a number of questions, which led us to a comprehensive analysis of mechanoreceptor and nociceptors. Do mechanoreceptors which transmit sensory input provided by sensorimotor activation (e.g. treadmill training) into the dorsal horn keep by these means nociceptor firing in check? Ablation of mechanosensory neurons in mice undergoing SCI would allow us to address this question. Specific genetic ablation of tropomyosin receptor kinase B (TrkB) expressing low threshold mechanoreceptors (LTMR) was previously investigated in a spared nerve injury model (5).

Is aberrant sprouting of nociceptor fibers in the dorsal horn really a key link explaining NP manifestation? Previously we could demonstrate that increased CGRP-positive peptidergic fiber sprouting in lamina III/IV below injury level following thoracic contusion SCI in mice can be reduced by sensorimotor activation (2, 3). Unfortunately, genetic ablation of CGRP-positive nociceptors was not successful (Armin Blesch, personal communication). Alternatively, C and Aδ nociceptive fibers were specifically ablated targeting sensory neuron specific (SNS; Nav1.8) expressing neurons, which allowed to probe their functional contribution to mechanical allodynia following PNS injury (6).

Can we identify potential drivers of SCI-induced NP in the DRG? As previously described, activity in nociceptors is likely the most important driver of NP in SCI (1). This assumption was primarily based upon DRG-derived nociceptor cultures taken from SCI animals, which indicated nociceptor hyperexcitability. While planning this study, such changes have not been reported yet in *ex vivo* skin-nerve preparations from below injury level, which are considered to be a better approximation towards pathophysiological events in the living animal compared to DRG cultures.

The relevance and comparability of animal SCI models in respect to the situation in human SCI subjects is very important. Of course, identical analyses are not applicable to human subjects for obvious reasons. However, quantitative sensory testing (QST), which stands for a battery of non-invasive psychophysical measures, allows to assess the function of the somatosensory system and - more specifically - hypersensitivity towards pain and mechanical stimuli (7), thus matching preclinical assessments of nociceptors (SNS^tdTomato^ transgenic mice for morphological analysis in the dorsal horn, electrophysiological properties of nociceptors *ex vivo* in wildtype SCI mice) and mechanoreceptors (ablation in TrkB^iDTR^ transgenic mice, electrophysiological properties of mechanoreceptors *ex vivo* in wildtype SCI mice).

The main aim of the present study was to comprehensively understand the initiation of NP signals and related structural rearrangements in the dorsal horn below injury level. In skin-nerve preparations from mice following thoracic contusion SCI, the evoked excitation level of mechanoreceptors and nociceptors was determined. Findings were correlated with non-invasive psychophysical measures (QST) reflecting the state of pain/touch hypersensitivity in SCI subjects up to 1 year after injury. Furthermore, genetic ablation of mechanoreceptors and nociceptors in respective transgenic mice was employed to investigate its relevance in pain behavior and structural rearrangement in the dorsal horn. Pre-SCI ablation of TrkB-expressing mechanoreceptors initially reduces mechanical allodynia but is not sustained. However, hyperexcitability was identified at 7 days post-SCI in nociceptors, not in mechanoreceptors, which in connection with rearranged peptidergic nociceptive fibers in lamina III/IV of the dorsal horn pave the way towards NP susceptibility and eventually NP manifestation.

## MATERIALS AND METHODS

### Animal subjects and experimental groups

Adult female C57BL/6J mice (8-12 weeks old at experimental initiation; wildtype JANVIER LABS) of varying genotypes subdivided into different experiments weighing between 20-25 g were used in this project. Animals were housed in groups of 4-5 mice per cage on a 12/12-hour light/dark cycle with access to water and food *ad libitum.* To ensure optimal conditions for housing and behavioral testing, the facility was controlled for temperature (22°C ± 1°C) and humidity (45-65 %) on a daily basis. All experiments were planned and conducted according to the PREPARE and ARRIVE guidelines (8, 9).

For experimental purposes and to ensure that the same investigator can perform behavioral testing, animals were divided in different cohorts for each experiment. Data from the different cohorts were combined in the end for statistical analysis.

### Spinal cord injury

All contusion surgeries were conducted with the Infinite Horizon Impact Device (IH-0400 Impactor; Precision Systems & Instrumentation, Lexington, KY, USA) adapted for the mouse spinal cord using a standard mouse steel-tipped impactor with a diameter of 1.3 mm (2, 3). Prior to surgery, mice were deeply anesthetized with an intraperitoneal (i.p.) injection using a cocktail (2.5mL/kg) containing ketamine (31.25 mg/kg), xylazine (1.58 mg/kg), and acepromazine (0.31 mg/kg). T9 laminectomy (corresponding to the T11 spinal level) was performed on mice of the injury group followed by a moderate contusion with a force of 50 kDyn. After contusion, the paravertebral muscle layers were sutured (4/0 terylene) and the skin incision was stapled. In sham mice, skin incisions and separation of paravertebral muscles to isolate the T9 vertebra was performed followed by suturing and stapling of the skin. Post-surgical care included analgesic medication, volume substitution, manual bladder voiding twice per day, and prophylactic antibiotic treatment to prevent bladder infection for 10-14 days.

All mouse experiments were conducted in accordance with the European Communities Council Directive (Directive 2010/63/EU amended by Regulation (EU) 2019/1010 and institutional guidelines) and approved by the local governing body (Regierungspräsidium Karlsruhe, Abteilung 3 - Landwirtschaft, Ländlicher Raum, Veterinär- und Lebensmittelwesen’, Germany (approval number: G-196/15).

### Transgenic animals

Crossing SNS^Cre^ (SCN10a promoter) with Rosa26^LSL-tdTomato^ reporter mice generates SNS^tdTomato^ mice in which peptidergic and non-peptidergic nociceptive neurons (Nav1.8^+^) can be traced by tdTomato. A total of 32 adult female SNS^tdTomato^ mice were assigned to either a sham operated control or SCI group and either analyzed 7 days post-injury (dpi) (SCI-7 dpi: n=4; Sham -7 dpi: n=3) or 21 days post-injury (SCI-21 dpi: n=16; Sham -21 dpi: n=9). Due to variation in offspring size and the inclusion of female mice only, behavioral and histological analysis for animals in which sprouting was evaluated 21 days post-injury was performed in three cohorts (numbers within parentheses indicate the group size for cohort 1, 2 and 3, respectively): SCI (n=4; n=3; n=9), Sham (n=3; n=2; n=4).

To investigate the contribution of nociceptive and mechanosensory neuronal subpopulations to SCI-induced below-level NP, heterozygous TrkB^CreERT2^::Avil^iDTR^ (TrkB^iDTR^) and SNS^Cre^::Rosa26^iDTR^ (SNS^iDTR^) mice (5, 6) in which either mechanosensory or nociceptive neurons can be specifically ablated have been used. To generate heterozygous TrkB^CreERT2^::Avil^iDTR^ (TrkB^iDTR^) animals, C57BL/6-Tg(TrkB-CreERT2)^1Phep^/Embph (TrkB^CreERT2^) mice were crossed to C57BL/6-Tg(Avil-^tm1-DTR^)^1Phep^/Embph (Avil^iDTR^) mice. Crossing SNS^Cre^ mice with C57BL/6-Gt(ROSA)26Sor^tm1(HBEGF)Awai^ mice (ROSA26^iDTR^) results in SNS^Cre^::Rosa26^iDTR^ (SNS^iDTR^) mice. Administration of tamoxifen (75 mg/kg i.p. for 5 consecutive days) into TrkB^iDTR^ mice specifically renders Avil-expressing TrkB^+^ sensory neurons susceptible to ablation by diphtheria toxin (DTX). Adult female SNS^iDTR^ (n=6) and TrkB^iDTR^ (n=19) mice were injected i.p with 40 μg/kg DTX (Sigma, D0564) with a second dose after 72 h. In TrkB^iDTR^ animals, DTX was administered 14 days after the last tamoxifen injection unless described otherwise. Control animals were injected i.p. with 100 μL saline twice in the same time frame. TrkB^iDTR^ animals were used to investigate the effect of ablation <u>after</u> SCI when mechanical allodynia has developed (SCI n=17; +Tam/+DTX n=9; +Tam/+saline n=6) or on development of mechanical allodynia (<u>before</u> SCI n=18; +Tam/+DTX n=10; +Tam/+saline n=8).

In SNS^iDTR^ mice it is possible to specifically ablate the Nav1.8^+^ cell population. However, diphtheria toxin-mediated ablation induced severe seizures (n=3) and even was lethal for three additional animals. Therefore, studies using these mice were discontinued.

### Behavioral Testing

Mice were habituated to the individual testing set-up for 2-3 days for at least 1.5 h per day prior to behavioral testing and for 30-60 min in each set-up on testing days. All behavioral testing was performed in awake, unrestrained mice by the same investigator blinded to group identity.

Basso Mouse Scale: Recovery of motor function was assessed using the Basso Mouse Scale (BMS) for locomotion as described before (2, 3). The motor function was scored 1 day post-injury to confirm an adequate injury and weekly starting 7 days post-injury. Each hind paw was scored individually and averaged to get a single score for each animal per testing day. A score of 0 reflects complete hindlimb paralysis without ankle movement, whereas a score of 9 indicates normal locomotion. The BMS was used to evaluate sensorimotor function to ensure proper recovery of motor function for sensory testing, which requires the animals to show weight support (BMS ≥ 3).

Mechanical sensitivity testing: To evaluate changes in mechanical sensitivity after injury and ablation of specific sensory neuronal subpopulations, von Frey hair filaments with ascending diameter and force (0.16 g, 0.4 g, 0.6 g, 1.4 g; Touch Test Sensory Evaluators; North Coast Medical, Gilroy, CA) were applied to the hindpaws of mice as described before (2, 3). For TrkB^iDTR^ animals, two additional filaments with lighter touch (0.04 g and 0.07 g) were included.

Starting with the smallest filament (0.16 g; 0.04 g for TrkB^iDTR^) the plantar surface of each hindpaw was stimulated as previously described resulting in 10 stimuli per filament per mouse (2, 3). To avoid stimulus-induced sensitization of the hind paw, up to 12 animals were tested in parallel and hind paws were stimulated in an alternating manner. Mechanical sensitivity towards each filament is defined as paw withdrawal frequency (number of positive responses divided by the number of stimuli) and expressed as the average for both hindpaws. Each animal was tested on 2 consecutive days before surgery and the mean was defined as the pre-operative baseline response rate.

To investigate nocifensive response duration (removal, licking and shaking) of a subcohort of wildtype mice (SCI n= 4, sham n=4), video recording of the response towards the 0.16 g and 1.4 g von Frey filament was performed pre-operatively and 7 days post-injury (iPhone XR, 1080p HD, 240 fps, slow-motion function, 4.16 ms/frame). Response duration was defined as the time until the hindpaw was placed down again.

Place Escape/Avoidance Paradigm: The place escape/avoidance paradigm (PEAP) is based on the assumption that animals escape and/or avoid an aversive mechanical stimulus. Animals have the active choice between a naturally preferred dark environment or pain relief in a more aversive light environment (10). Here, it was employed to address the aversive quality of pain and thereby supraspinal processing of nociception as previously described (3). In short, animals were placed in the middle of the chamber and the time spent on the dark side of the chamber was measured. An initial 10 min exploration phase (baseline) without any stimulus was followed by a 15 min testing phase (3× 5-min blocks) in which the hindpaws were stimulated using a 0.16 g von Frey filament every 15 sec when the mouse was present on the dark side of the chamber. The left and right hindpaws were stimulated in an alternating manner.

Thermal sensitivity testing: Thermal sensitivity was assessed according to Hargreaves’ method using the plantar test (Hargreaves Apparatus; Ugo Basile) as previously described (2, 3). In short, an infrared laser beam (190 ± 1 mW/cm^2^) was presented to the plantar surface of the hindpaws and the time (sec) until withdrawal was recorded resulting in the response latency to the heat stimulus (11). To avoid tissue damage, a 15-sec cut-off time, as well as a delay of at least 1 min between two trials for the same paw and mice, was introduced. In each animal, the mean of four trials for the left and right hindpaws was used to express thermal sensitivity and testing was always performed after mechanical sensitivity was assessed. Similar to von Frey testing, the mean latency of 2 consecutive testing days before surgery or any other intervention determined the baseline of the animals (pre-operatively; pre-intervention).

### Tissue Processing and morphological analysis

All animals were euthanized with an overdose of anesthesia cocktail (ketamine, xylazine, and acepromazine) and transcardially perfused with 0.9 % saline followed by 4 % paraformaldehyde (PFA) in 0.1 M phosphate buffer (2). The spinal column was dissected, post-fixed in 4 % PFA for 1 h at room temperature, and briefly rinsed with ddH_2_O followed by a 5 min washing in 0.1 M phosphate buffer before the tissue was cryoprotected in 30 % sucrose in 0.1 M phosphate buffer at 4°C. The lesion site (T11; if applicable), the L4-L6 spinal cord, as well as L4-L6 DRG were identified using anatomical landmarks (12), embedded in Tissue-Tek® O.C.T compound (Sakura Finetek, Germany) followed by serial sectioning (spinal cord, 16 and 25 µm coronal; DRG, 10 µm) directly on to slides as previously described (2, 3).

Eriochrome Cyanine staining: The lesion size of injured animals was analyzed by quantifying the spared tissue at the lesion epicenter. Two consecutive series were stained for myelin using Eriochrome-Cyanine (EC) as previously described (2, 3). Tissue sparing at the lesion epicenter (smallest area of spared white matter) was quantified by outlining areas with light and dark blue staining and expressed as the percent of cross-sectional area calculated by dividing the white matter area by the total cross-sectional area. Imaging and analysis was performed blinded to group identity using ImageJ.

Immunohistochemistry: Immunohistochemical labeling was performed on slides with serial sections of the lumbar spinal cord (L4-L6) and the corresponding L4-L6 DRG. Slides were dried for 30 min at room temperature, encircled with a liquid blocker and rinsed in 0.1 M Tris-buffered saline (TBS). Sections were blocked and permeabilized by incubating in TBS/0.25 % Triton TX-100/5 % donkey serum followed by incubation with primary antibodies diluted in TBS/ 0.25 % Triton TX-100/1 % donkey serum overnight at 4°C. On day 2, sections were incubated with secondary antibodies and 4’,6-diamidino-2-phenylindole (DAPI), rinsed, dried and cover-slipped with Fluoromount G (Southern Biotechnology Associates, Birmingham, AL) (3). For each animal, images of the left and right dorsal horn of three labeled sections with an intersection distance ranging from 700-1400 µm were taken at 20 × magnification with the same resolution, lens aperture, and exposure time using a XC30 camera mounted on an Olympus BX53 microscope with epifluorescence illumination and appropriate filter cubes. Lumbar DRGs (L4-L6) were imaged bilaterally from each animal at 10 × magnification using the above-described set-up, and images from three sections per DRG (140 µm apart) were analyzed.

The following primary antibodies were used: rabbit anti-CGRP (Immunostar, 24112; 1:200), rabbit anti-protein kinase Cγ (PKCγ [C19], Santa Cruz Biotechnology, SC-211; 1:200), guinea pig anti-NeuN (Millipore, ABN90; 1:2000) and goat anti-human HB-EGF (R&D Systems, AF-259-NA; 1:40). Secondary antibodies were Alexa Fluor 488 and Alexa Fluor 594 donkey anti-rabbit and donkey anti-goat (all Life Technologies; 1:300), Alexa Fluor 594-conjugated streptavidin (Jackson ImmunoResearch; 1:300), Alexa Fluor 488 donkey anti-guinea pig (Jackson ImmunoResearch; 1:300).

Quantification of labeling density in the spinal cord: The termination pattern of peptidergic-and non-peptidergic fiber was analyzed in lamina III-IV of the dorsal horn as previously described (3). In short, an analysis box was placed in the center of the dorsal horn adjoining the ventral border of lamina IIi which was identified by PKCγ-labeling. The dorsoventral extent of lamina IIi was used as a reference for the dimensions of the region of interest (ROI) analysis box. Images were processed by setting a labeling threshold minimizing background and accurately reflecting CGRP- and Tomato-positive fiber labeling. Labeling density was expressed as percentage of positive labeling within the analysis box and the values for three sections per animal were averaged. Quantification of labeling density was performed blinded to group identity using ImageJ.

Quantification of ablated DRG neurons: In TrkB^iDTR^ L4-L6 DRG quantification of HB EGF^+^ in NeuN^+^ neurons was used to evaluate the proportion of neurons expressing DTR (DTR^+^/NeuN^+^).

### Ex vivo skin nerve recordings

Skin-nerve recordings were performed on 10-12 week-old SCI or Sham mice killed with CO_2_ followed by cervical dislocation. The hindpaw skin was dissected free together with the sural nerve and was placed in an organ bath chamber that was perfused with 32°C-warm synthetic interstitial fluid (SIF) consisting of 108 mM NaCl, 3.5 mM KCl, 0.7 mM MgSO4, 26 mM NaHCO3, 1.7 mM NaH2PO4, 1.5 mM CaCl2, 9.5 mM sodium gluconate, 5.5 mM glucose and 7.5 mM sucrose at a pH of 7.4. The sural nerve was led through a small hole into the adjacent mineral oil filled recording chamber. The nerve was teased into thin bundles that were laid on a silver wire recording electrode connected to a differential amplifier (Digitimer, modules NL104, NL125/NL126) and nerve fiber activity was recorded with the Powerlab 2 4SP system and Labchart 7.1 software (AD Instruments). The receptive fields of single nerve fibers were located by manually probing the skin using von Frey hair filaments. To distinguish between Aβ-fibers, Aδ-fibers and C-fibers, action potentials were evoked by electrical stimulation of the receptive fields and the axonal conduction velocities were calculated by dividing the distance between the stimulation electrode and the recording electrode by the delay between the onset of electrical stimulation and the arrival of the action potential at the recording electrode. Nerve fibers with a conduction velocity greater than 10 m/s were classified as Aβ-fibers, fibers with a CV between 1 and 10 m/s as Aδ/fibers and fibers with a CV < 1m/s as C-fibers. To further distinguish between nociceptors and mechanoreceptors, the mechanical activation thresholds were determined with von Frey hair filaments. Finally, to determine the action potential firing rates and adaptation patterns at different stimulus strengths, the receptive fields were stimulated with ramp-and-hold stimuli applied with a linear piezo actuator (Nanomotor®, Kleindiek) equipped with a force measurement system (FMS-LS, Kleindiek) to measure the exact force of the applied stimulus.

### Human SCI study

Included participants (n=17) were identified and recruited from July 2016 through June 2021 by convenient sampling. All of them provided written informed consent. Six individuals did not complete the final timepoint. The human SCI study was conducted within the framework of the European Multicenter Study about Spinal Cord Injury at the SCI center, Heidelberg University Hospital (*ClinicalTrials.gov* register-no. NCT01571531; https://emsci.org). The study protocol was approved by the local ethical review board (S-188/2003).

Sensory function was assessed in SCI subjects by both the International Standards for Neurological Classification of SCI (ISNCSCI) and highly standardized Quantitative Sensory Testing (QST) (13, 14). QST represents a comprehensive protocol for assessing the sensory system and is reported to give indication of sensory tract integrity concerning both the peripheral and the central nervous system. For the classification of existing pain problems in individuals with SCI all participants received the clinical assessment EMSCI Pain Assessment Form (EPAF) that is in accordance with international guidelines and recommendations for the clinically documentation and assessment of pain in SCI (15). All indivuduals received this assessment in the three mentioned time points (1, 3 and 12 months).

In consideration of the methodological approach for the animal model, special focus was set on the evaluation of the mechanical detection threshold (MDT) representing a test for Aβ-fibers (mechanoreceptors) and the mechanical pain threshold (MPT) to function as indicator for Aδ-fibers (nociceptors) (16–18). While MDT was performed via modified von Frey hair filaments (contact area 0.5 mm in diameter; Optihair2-Set, Marstock Nervetest, Germany) applying forces from 0.25-512 mN, starting with 8 mN, MPT was assessed by means of (weighted) pin prick examination (contact area 0.25 mm in diameter) applying forces from 8 to 512 mN, starting with 8mN (18).

Dedicated areas being tested as to alterations of sensory function were the dermatomes L4 (medial right shin, mid-way between knee and ankle) and L5 (dorsum of the foot, third metatarsal phalangeal joint). The rationale for choosing L4/5 was based on related literature reporting a frequent localization of NP in these regions (19–22).

Evaluation and analysis of QST data was done based on standard values (respecting the tested areas of the body, gender, and age) as provided by the German Research Network on Neuropathic Pain (DFNS) by means of a related database (23, 24).

### Statistical Analysis

Behavioral results were analyzed by repeated measures (RM) Two-way analysis of variance (ANOVA) to reveal overall group and timepoint differences, without assumption of sphericity (Geisser-Greenhouse corrections). Significant group differences were followed by Sidak pairwise comparisons. Significant timepoint differences between baseline vs post-SCI or baseline vs. post-DTX were followed by *post hoc* Fisher’s least significant difference (FLSD) test.

For experiments in which transgenic animals have been used, the lesion size, fiber density in the lumbar spinal cord and changes in DRG neurons (ablated neurons) were only compared between saline injected control SCI and ablated SCI group and consequently analyzed by unpaired t-test. For tdTomato-labeled transgenic animals with multiple timepoint groups, One-way ANOVA with *post hoc* FLSD was used to analyze lesion size and labeling density.

Electrophysiological data was analyzed with multiple Chi-Squared tests and multiple Mann-Whitney tests.

All data are presented as mean ± standard deviation (SD), except for skin-nerve preparation data which is presented as mean ± standard error of mean (SEM). Statistical analysis was done using Prism 9 software (GraphPad Software Inc., La Jolla, CA) with an alpha level of 0.05 for significance.

QST data was processed and analyzed using the software ‘eQUISTA’ (Casquar GmbH, Bochum, Germany). Accordingly, results are expressed as Z-transformed standard values (‘Z-scores’). Values greater than 2 and lower than -2, indicating 2 standard deviations from healthy matched controls findings, meaning hyper- or hyposensitivity to the respective stimulus (14, 25).

## RESULTS

### Pre-injury ablation of TrkB-expressing mechanosensory neurons reduces NP behavior following mouse contusion SCI

We hypothesized that low-threshold mechanoreceptors (LTMRs), responsible for light touch, mediate beneficial sensory input provided by natural locomotor activity or in an enhanced setting such as treadmill training (4, 26–28). Therefore, we investigated the contribution of TrkB-expressing LTMRs (identified as Aδ-LTMRs, D-hair, and rapidly adapting Aβ mechanoreceptors, Aβ-RAMs) to SCI-induced mechanical allodynia by ablating TrkB-expressing LTMRs in TrkB^CreERT2^:Avil^iDTR^ (TrkB^DTR^) mice.

First, to determine the contribution of TrkB-expressing sensory neurons to established signs of SCI-induced neuropathic pain, respective neurons were genetically ablated 14 days following thoracic contusion SCI using diphtheria toxin (DTX), when mechanical allodynia and thermal hyperalgesia have already developed (Fig.1A). Prior to SCI, tamoxifen administration did not affect mechanical and thermal sensitivity (data not shown) and as expected all SCI animals developed mechanical allodynia and thermal hyperalgesia at 7 days post-injury (Fig. 1E-G, I, Two-way RM ANOVA, time differences p<0.0001). Genetic ablation resulted in a 97% reduction in TrkB^+^/DTR^+^ neurons (Fig. 1B-D; t-test p<0.0001 DTX vs. saline). Despite the high ablation efficiency, no significant effect of post-SCI ablation on mechanical hypersensitivity was observed. Statistical analysis indicated no overall group differences comparing the response rates for the small diameter von Frey hair filaments (0.04 g - 0.16 g) between saline-injected control and ablated SCI animals (Fig. 1 E-G, Two-way RM ANOVA Group: 0.04 g, p=0.54; 0.07 g, p=0.27; 0.16, g p=0.06). In line with this observation, supraspinal processing assessed by the place escape/avoidance paradigm (PEAP) also showed no difference between groups in response to the 0.16 g filament (Fig. 1 H, Two-way RM ANOVA p=0.23 for group differences).

**Figure 1:**
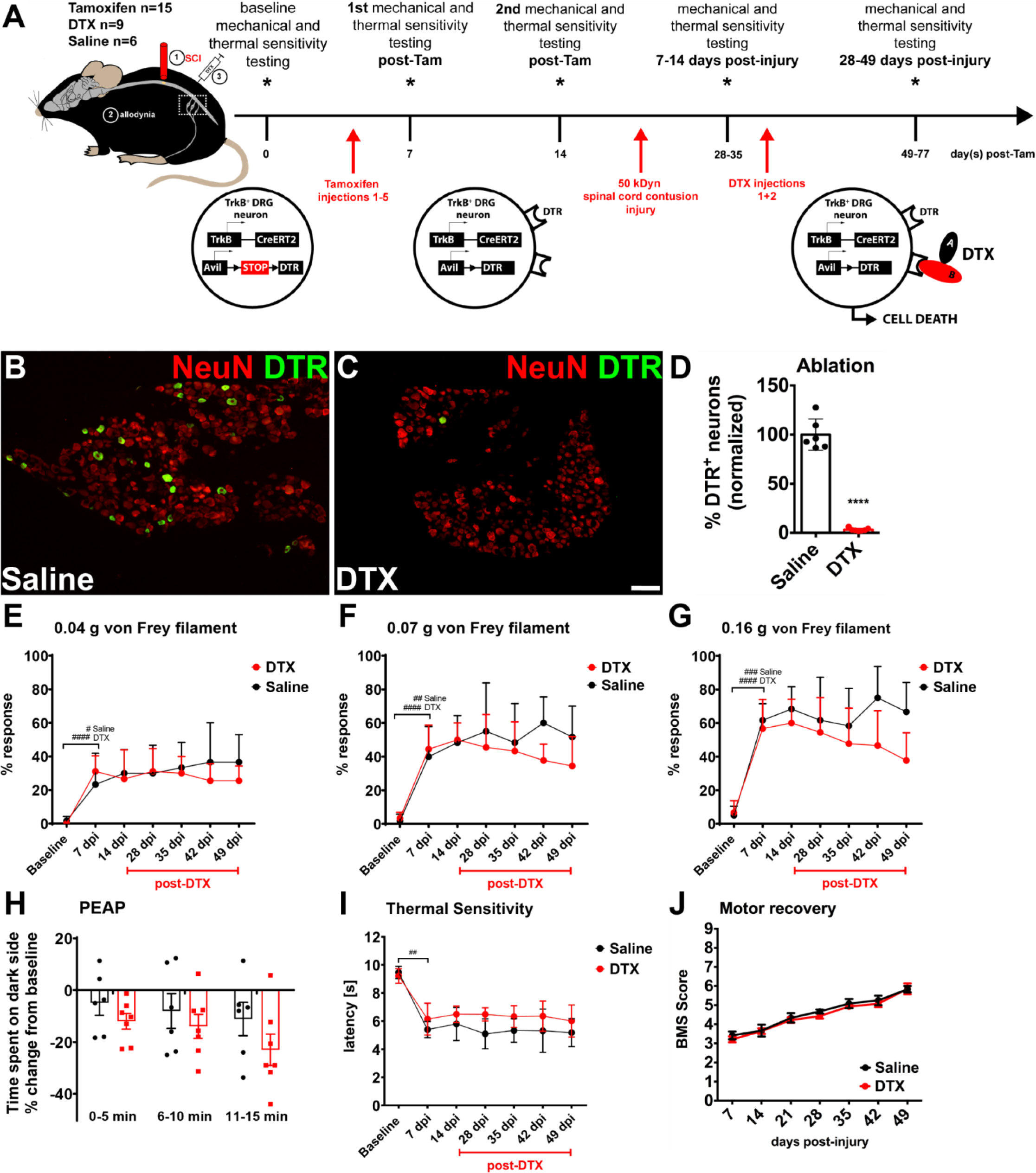
Post-SCI ablation of TrkB expressing mechanoreceptors. (**A**) Schematic of Experimental design. (**B**) Representative lumbar (L4-L6) DRG neurons of saline-treated control animals stained for HB-EGF shows robust expression of DTR in a subset of sensory neurons. (**C**) In ablated TrkB^DTR^ animals, DTR expression is almost completely absent. (scale bar, 100 μm) (**D**) Quantification (t-test p< 0.0001; DTX, n=7; Saline n=6, L4-L6 bilaterally). (**E-G**) SCI induced mechanical hypersensitivity to small-diameter light touch eliciting von Frey hair filaments (0.04 g, 0.07 g, 0.16 g, Two-way RM ANOVA, time differences p<0,0001, baseline vs 7dpi FLSD # p<0.05, ## p<0.01, ### p<0.001, #### p<0.0001), but DTX-mediated ablation of TrkB+ sensory neurons (28-49 dpi) did not change SCI-induced mechanical hypersensitivity. (**H**) The Place Escape Avoidance Paradigm (PEAP, 0.16 g) performed at the end of the study shows that both groups spent more time on the light side of the chamber (Two-way RM ANOVA p=0.23 for group differences). (**I**) Injured TrkB^DTR^ animals of both groups develop thermal hypersensitivity assessed by the Hargreaves test (Two-way RM ANOVA, baseline vs 7dpi p<0,0001, FLSD ## p<0.01, group differences p=0.02, no significant Sidak pairwise comparisons). (**J**) Except for the SCI-induced motor deficits scored by the Basso Mouse Scale (BMS) no further motor deficits are observed following post-SCI ablation of TrkB^+^ sensory neurons. Mean ± SD for all graphs.

Post-SCI ablation did not affect response rates measured using larger diameter innocuous von Frey hair filaments (Suppl. Fig. 1 A, 0.4 g: Two-way RM ANOVA p=0.17), but did have significant group differences for noxious stimuli (0.6 g and 1.4 g) with inconsistent or nonsignificant pairwise comparisons at each timepoint (Suppl. Fig. 1 B-C; Two-way RM ANOVA, group differences 0.6 g, p=0.04; 1.4 g, p=0.01). TrkB-expressing LTMR ablation did not affect motor recovery (Fig. 1 J). Following SCI, significant thermal hypersensitivity (Hargreaves’ method) was observed for all SCI animals irrespective of ablation (Fig. 1 I, Two-way RM ANOVA, baseline vs 7 dpi p<0,0001, group differences p=0.02, no significant Sidak pairwise comparisons). No significant differences in white matter sparing (lesion size) were observed between saline-injected controls and ablated SCI mice (Suppl. Fig. 2 A; t-test p=0.52). Taken together, these results suggest that TrkB-expressing sensory neurons do not play a role in already developed and stably present mechanical allodynia following SCI.

Second, to determine the role of TrkB-expressing sensory neurons in the initiation of SCI-induced neuropathic pain, neurons were ablated prior to SCI. This resulted in an effective reduction of >90 % of DTR^+^ neurons in lumbar DRG (L4-L6) (Fig. 2 B-D; t-test p<0.0001 DTX vs. saline control group). Without injury no differences in mechanical or thermal sensitivity were observed (Fig. 2 E-G, I, Suppl. Fig. 1 D-F). All SCI animals of the saline-injected control group showed mechanical hypersensitivity at 7 days post-injury compared to pre-SCI values using small-diameter von Frey filaments (Fig. 2 E-G). Interestingly, a significant group effect was observed (Fig. 2 E-G; Two-way RM ANOVA group differences: 0.04 g, p=0.004; 0.07 g, p=0.008; 0.16 g, p=0.013). However, TrkB-ablated SCI animals (DTX group) showed a lower response rate than the control group only at 21 days post-injury indicating a small but significant effect of pre-injury ablation of TrkB-expressing sensory neurons on SCI-induced mechanical allodynia. In line with the lack of group differences to the 0.16 g von Frey hair filament at the end of the experiment, the PEAP indicated SCI-induced mechanical allodynia in both groups (Fig. 2 H, Two-way RM ANOVA group differences p=0.49). Pre-SCI ablation did not affect response rates measured using larger diameter (0.4 g - 1.4 g) von Frey hair filaments (Suppl. Fig. 1 D-F, Two-way RM ANOVA, group differences 0.4 g, p=0.11; 0.6 g, p=0.11; 1.4 g, p=0.13). Both, thermal sensitivity (Fig. 2 I, p=0.14) and motor recovery (Fig. 2 J) were not influenced. Additionally, differences in sensory behavior of saline-injected and ablated SCI animals cannot be attributed to difference in white matter sparing (lesion size) (Suppl. Fig. 2 B, t-test p=0.69). Taken together, only pre-injury – not post-injury - ablation of TrkB-expressing mechanosensory neurons reduces SCI-induced mechanical allodynia.

**Figure 2:**
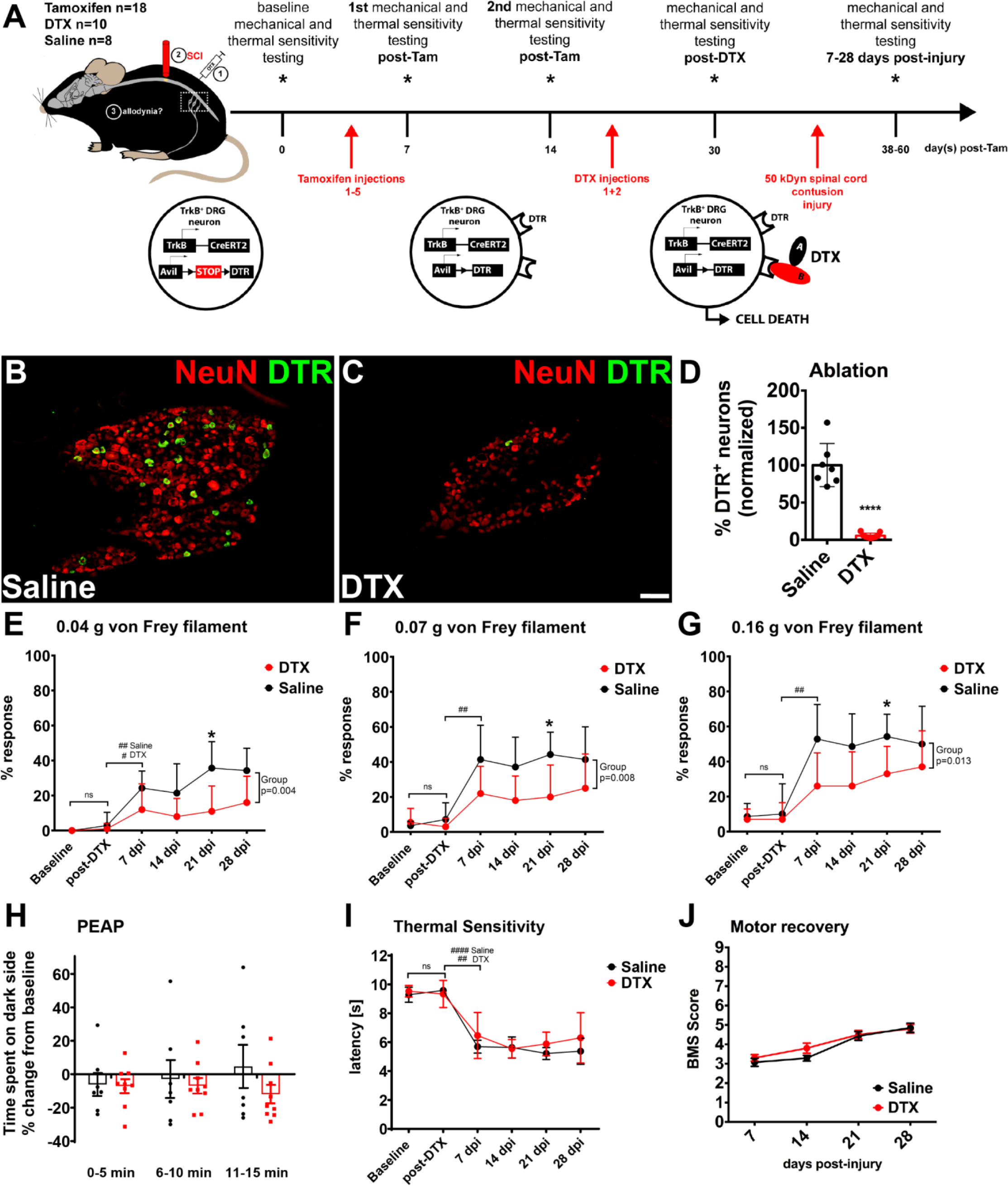
Pre-SCI ablation of TrkB-expressing mechanoreceptors. (**A**) Schematic of Experimental Design. (**B**) Saline-injected control animals show robust DTR expression in lumbar (L4-L6) DRG neurons. Compared to control animals, DTR+ neurons are almost completely ablated in DTX-injected animals (**C**) (scale bar, 100 μm). (**D**) Quantification (t-test p< 0.0001; DTX, n=9; Saline n=7, L4-L6 bilaterally). (**E-G**) SCI-induced mechanical hypersensitivity to small-diameter filaments (0.04 g, 0.07 g and 0.16 g, Two-way RM ANOVA, time differences p<0,0001, post-DTX vs 7 dpi FLSD # p<0.05, ## p<0.01, no significant (ns) difference between baseline and post-DTX) is significantly altered in ablated animals, but inconsistently (Two-way RM ANOVA group differences: 0.04 g, p=0.004; 0.07 g, p=0.008; 0.16 g, p=0013, Sidak pairwise comparisons * p <0.05). The response rate towards the 0.16 g von Frey hair filament (**G**) of ablated animals is not significantly different from control animals anymore at the end of the experiment 28 dpi. (**H**) This is confirmed by supraspinal processing of nociception using the PEAP (0.16 g) where both groups equally spent more time on the light side of the box. (**I**) DTX treatment does not affect pre- or post-injury thermal sensitivity, but SCI did induce thermal sensitivity in both groups (Two-way RM ANOVA, time differences p<0,0001, post-DTX vs 7dpi FLSD ## p<0.01, #### p<0.0001). (**J**) SCI induces significant motor deficits in all animals scored by the Basso Mouse Scale (BMS) and ablation of TrkB+ sensory neurons does not induce further motor deficits. Mean ± SD for all graphs.

Regardless of the ablation timing (post- or pre-SCI), CGRP-density in laminae III-IV of the lumbar dorsal horn was not reduced in respective transgenic SCI mice compared to the non-ablated SCI control group (Fig. 3 A-F). Therefore, SCI-induced maladaptive structural changes regarding sprouting of CGRP^-^expressing fibers were not influenced by ablation of TrkB-expressing sensory neurons and existed while mechanical allodynia was present.

**Figure 3:**
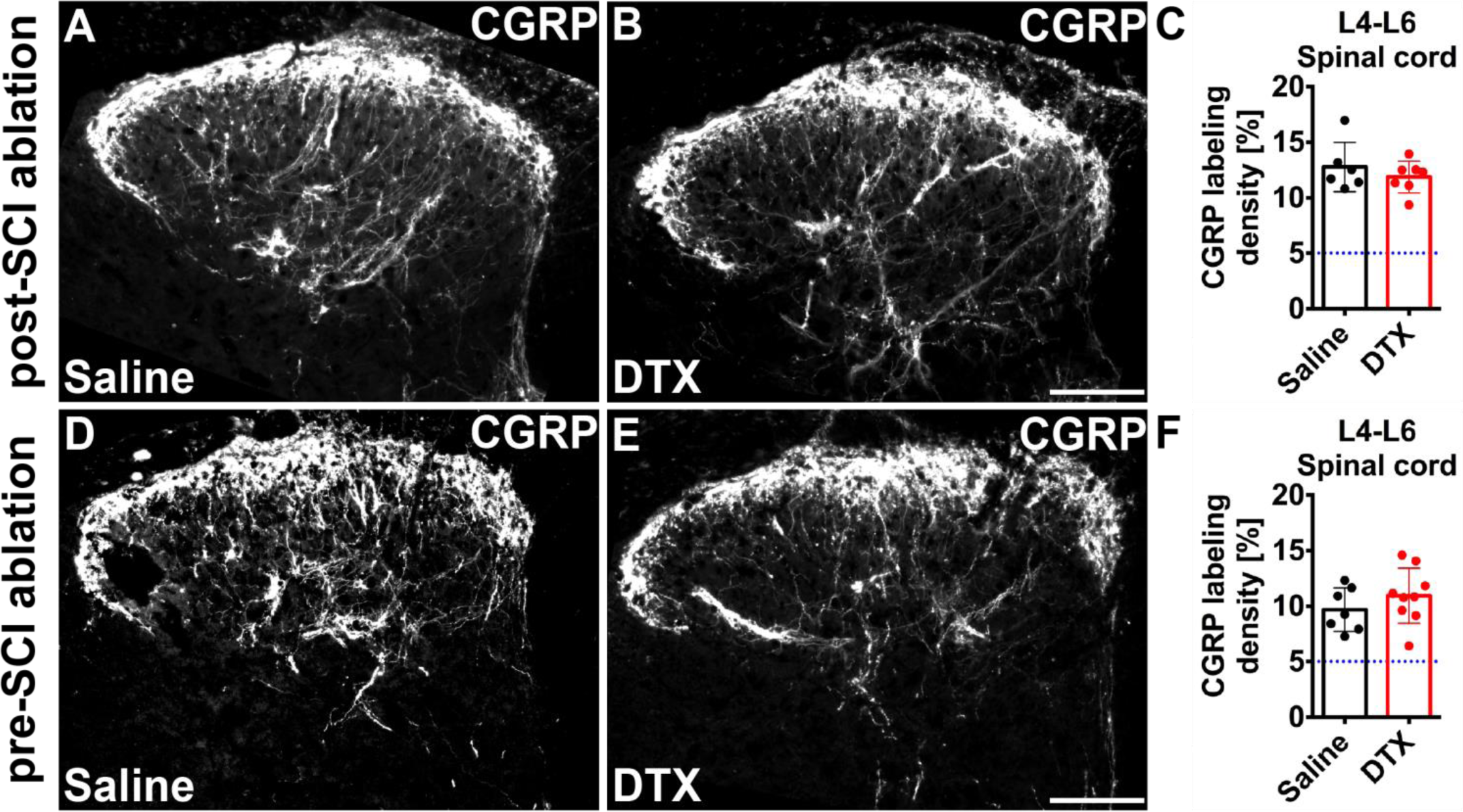
Peptidergic nociceptive fiber changes in the dorsal horn following TrkB-expressing mechanoreceptor ablation. Lumbar (L4-L6) spinal cord sections of injured saline control (**A, D**) and DTX-treated injured (**B, E**) TrkB^DTR^ mice stained for CGRP (scale bar, 100 μm). Representative images indicate the SCI-induced increase in CGRP-labeling density in deeper laminae (III-IV) of the spinal dorsal horn in all groups. (**C, F**) Quantification of the CGRP-labeling density in laminae III-IV of the lumbar (L4-L6) dorsal horn shows no difference comparing the injured saline-injected control group with injured DTX-injected animals but a significant increase compared to sham animals of Figure 4 (dotted line). (Mean ± SD) (post-SCI; Saline n=6; DTX n=7) (pre-SCI; Saline n=7; DTX n=9)

### Sprouting of nociceptive fibers parallels the development of SCI-induced mechanical allodynia

To determine the distribution and sprouting pattern of all nociceptive (peptidergic and non-peptidergic) neurons as opposed to only CGRP expressing peptidergic nociceptors, the temporal dynamics of nociceptive fiber sprouting in response to SCI and its relation to the development of NP was investigated. We therefore took advantage of a previously described sensory neuron specific (SNS; Na_v_1.8) Cre-driver line showing recombination in all nociceptive neurons of the DRG in combination with additional CGRP staining. This labeling and staining combination ensures that the observed SCI-induced structural changes are not specifically Na_v_1.8- or CGRP-dependent (Suppl Fig. 3).

The previously established SNS^tdTomato^ mice (6, 29) were examined for nociceptive fiber sprouting (3, 5, 7 and 21 days post-injury) and NP behavior before and 7, 14 and 21 days post-injury. These timepoints were chosen based on motor deficits as a consequence of SCI allowing the assessment of sensory changes only after weight support is re-gained (7 days post-injury; BMS 3-4), whereas at 21 days post-injury NP is fully established (2, 3).

As expected, following a moderate T11 spinal cord contusion injury mice developed mechanical allodynia (0.16g von Frey hair filament) as early as 7 days post-injury that persisted through 21 days post-injury (Fig. 4 A, Two-way RM ANOVA, group differences p=0.02). Hypoalgesia to noxious stimuli (0.6 g and 1.4 g von Frey hair filaments) developed and persisted as well (Fig. 4 A). Further, thermal hyperalgesia (Hargreaves’ method) started as early as 7 days post-injury and persisted through 21 days post-injury (Suppl. Fig. 4 A-B). All SCI animals were confirmed to have significant hindlimb motor deficits by BMS at all timepoints compared with Sham animals (Suppl. Fig. 4 C). In line with the motor deficits, quantification of white matter sparing indicated greater than 50% loss of white matter and gray matter damage at the lesion epicenter (Suppl. Fig. 4 D-E).

**Figure 4:**
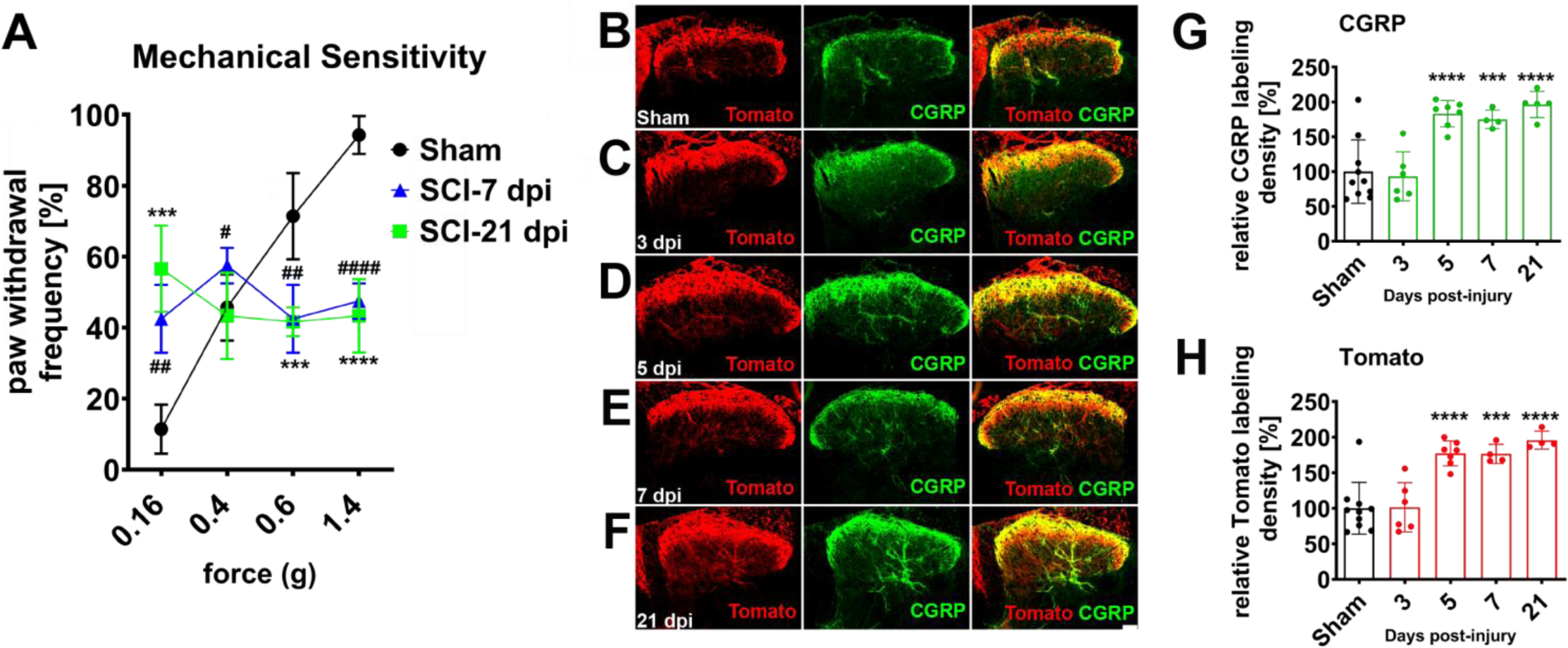
Nociceptive fiber changes in the dorsal horn. (**A**) SCI animals develop (7 dpi) and maintain (21 dpi) mechanical allodynia and hypoalgesia (Two-way RM ANOVA, Sidak pairwise comparisons SCI-7 dpi vs Sham: ## p<0.01, #### p<0.0001, SCI-21 dpi vs Sham: *** p<0.001, **** p<0.0001). Nociceptive plasticity into deeper laminae (III-IV) is shown with SNS^tdTomato+^ mice (**B**) Sham, (**C**) 3 dpi, (**D**) 5 dpi, (**E**) 7 dpi, (**F**) 21 dpi with both Tomato (**G**) and CGRP (**H**) staining quantified. We observe here that sprouting occurs as early as 5 dpi which is in line with the development of mechanical allodynia (One-way ANOVA, FLSD *** p<0.001, **** p<0.0001). Mean ± SD

To understand if SCI-induced aberrant structural changes arise prior to and may be responsible for the development of NP, nociceptive fiber sprouting was compared at 3, 5, 7 and 21 days post-injury. While at 3 days post-injury, no difference was observed comparing Sham and SCI mice (Fig. 4 B, C, G-H), at 5 days post-injury tdTomato- and CGRP-labeling density in laminae III-IV of the lumbar (L4-L6) dorsal horn was increased by approximately 70% in the SCI group and persisted through 21 days post-injury (Fig. 4 B, D-H One-way ANOVA p<0.0001; 5, 7 and 21 dpi Fischer least significant difference, FLSD p≤0.0001).

To further shed light on nociceptive neurons contributing to SCI-induced pain behavior, nociceptive Na_v_1.8-expressing neurons were genetically ablated in SNS^Cre^::R26^iDTR^ (SNS^iDTR^) mice (Suppl. Fig. 5). However, ablation of nociceptive neurons through intraperitoneal injection of DTX in SNS^iDTR^ mice induced seizures in some animals (n=3), which had to be sacrificed, while others died within hours (n=3).

### Experimental contusion SCI induces nociceptor, but not mechanoreceptor, hyperexcitability

To identify the key neuronal population (nociceptors or mechanoreceptors) driving nociceptive fiber sprouting in the dorsal horn and mechanical allodynia as correlate of NP behavior, *ex vivo* skin-nerve preparations were investigated.

Single-unit teased fiber recordings from *ex-vivo* hindpaw skin-nerve preparations revealed that C- and Aδ-fiber nociceptive sensory afferents displayed decreased mechanical activation thresholds 7 days post-injury (Fig. 5 A-E). Thus, a significantly higher proportion of C-fiber nociceptors responded to forces smaller than 125 mN (Fig. 5 E, multiple Chi-Squared tests). Moreover, C-fiber nociceptors fired significantly more action potentials in response to mechanical ramp-and-hold stimuli (Fig. 5 F, multiple Mann-Whitney tests) and exhibited continuous action potential firing after removal of the stimulus (Fig. 5 G-I, multiple Mann-Whitney tests). These afterdischarges were also observed in Aδ-fiber nociceptors (Fig. 5 G and H). Additionally, a significantly higher proportion of Aδ-fiber nociceptors responded to forces at 17.5 and 22.5 mN (Fig. 5 C, multiple Chi-squared tests); however, they did not fire more action potentials in response to mechanical ramp-and-hold stimuli (Fig. 5 D, multiple Mann-Whitney tests). Most importantly, no significant changes in the mechanical activation thresholds or the action potential firing rates were observed after SCI in the three major classes of low-threshold mechanoreceptors (RA Aβ-LTMRs, SA Aβ-LTMRs, DH Aδ-LTMRs, Fig. 5J–M). Therefore, a midthoracic contusion SCI induces hyperexcitability of peripheral nociceptive fibers but not mechanoreceptors of the hindpaw.

**Figure 5:**
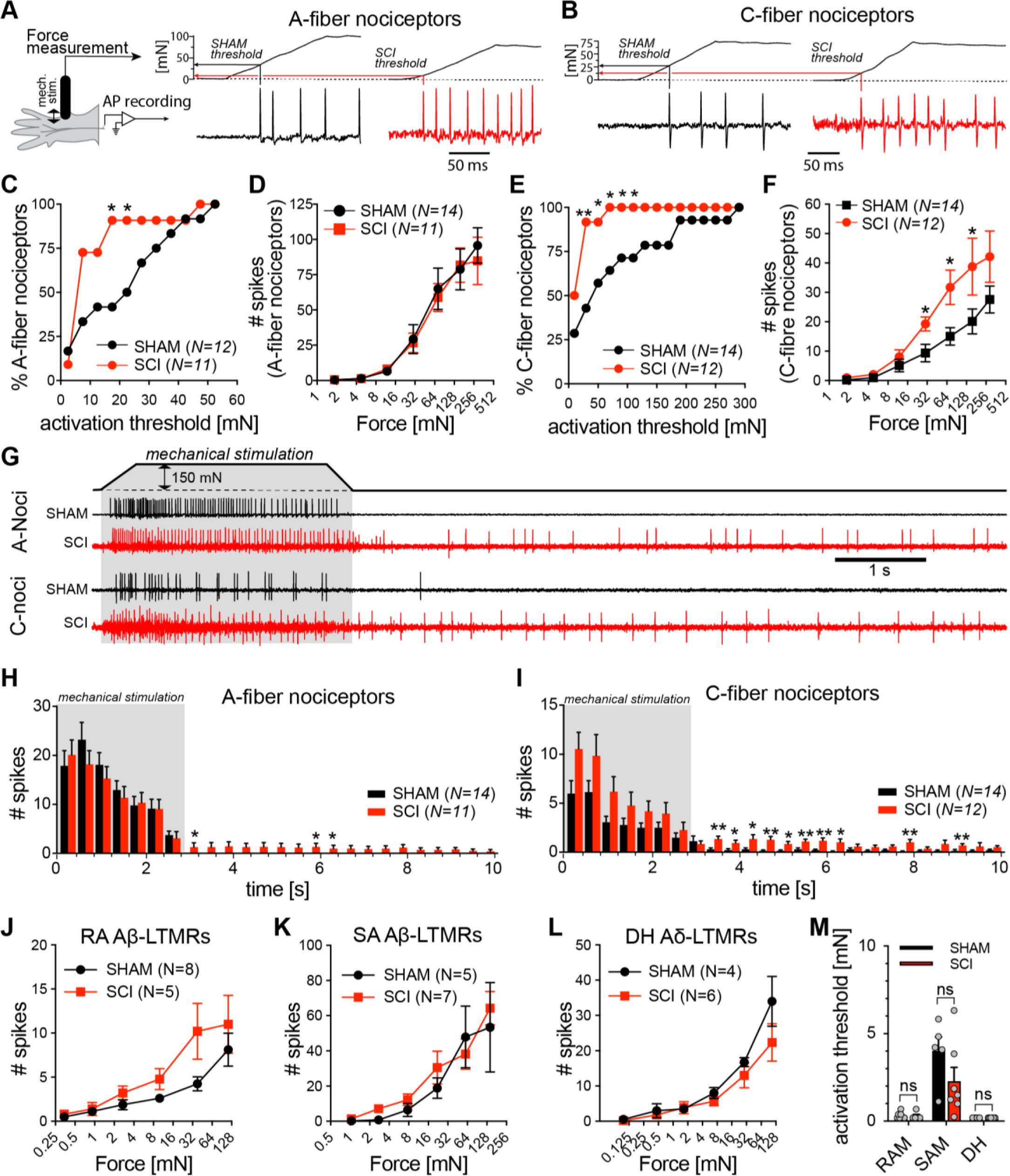
Mechanoreceptor and nociceptor activity after SCI. Single-unit teased fiber recordings from hindpaw glabrous skin-nerve preparations show A-fiber (**A**) and C-fiber nociceptors (**B**) have lowered activation thresholds 7 dpi T11 (50 kDyn) SCI in mice. This is quantitatively shown in (**C**) for A-fiber and (**E**) for C-fiber nociceptors responding to forces smaller than 125mN (multiple Chi-squared tests). While following SCI A-fibers (**D**) do not fire more frequently, C-fibers (**F**) significantly fire more frequently in response to mechanical ramp-and-hold stimuli. Example traces exhibit continuous firing following removal of stimulus (150mN denoted in gray area) in the SCI group (**G**). This is quantitatively shown for A-fiber (**H**) and C-fiber (**I**) nociceptors (multiple Mann-Whitney tests). The firing of RA-Aβ-LTMRs (**J**), SA-Aβ-LTMRs (**K**) and DH-Aδ-LTMRs (**L**) was not significantly altered by SCI. Additionally, LTMR activation thresholds were not altered by SCI (**M**). Mean ± SEM

Does prolonged nocifensive behavior such as removal, licking and shaking correlate with the extension of the firing past stimulus removal? Indeed, almost a doubling of the response duration of the SCI mice to the 0.16 g filament was observed at 7 days post-injury (Fig. 6 A, t-test p=0.03). Also the response rate for these mice was significantly increased 7 days post-injury (Fig. 6 B, Two-way RM ANOVA, group differences, p=0.03). Therefore, the continued firing observed in nociceptors appears to be behaviorally expressed by prolonged nocifensive behavior in SCI mice.

**Figure 6:**
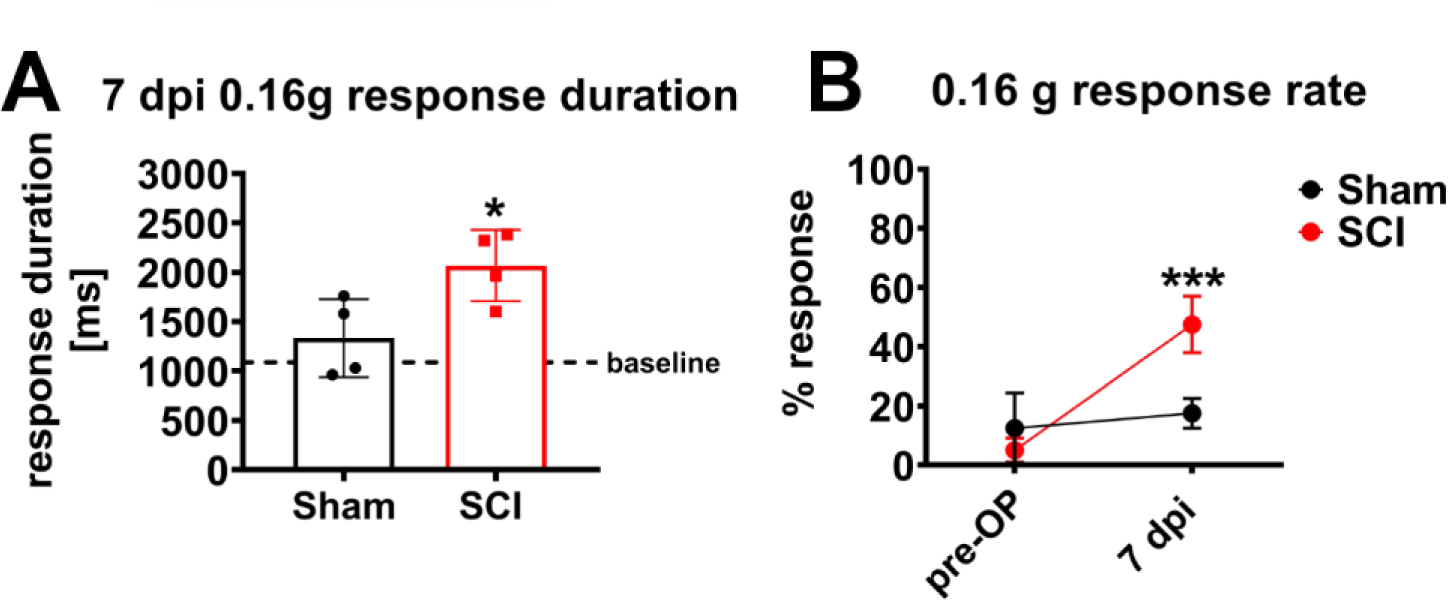
Prolonged nocifensive behavior following experimental SCI. (**A**) T11 SCI mice showed prolonged nocifensive behavior (licking, shaking and holding) compared to Sham mice when responding to 0.16g von Frey hair filaments 7dpi (slow-motion video analysis, unpaired two-tailed t-test, * p<0.05). (**B**) The response rate of SCI rate was significantly higher than Sham mice at 7dpi with 0.16g von Frey hair filament. (Sham n=4, SCI n=4, Two-way RM ANOVA, group difference p=0.03, Sidak pairwise comparison *** p<0.001)

### Nociceptor hypersensitivity is observed in SCI subjects up to one year post-injury

The best approximation towards the relevance in nociceptors versus mechanoreceptors in initiating/maintaining hyperexcitability and NP presentation, quantitative sensory testing (QST) was employed. QST was performed 1, 3 and 12 months post-injury in 17 conveniently sampled individuals with SCI varying in age (18 – 75 years), sex (male, n=13; female, n=4), cause of injury (traumatic, n=11; ischemic, n=5; or hemorrhagic, n=1), initial classification according to the ASIA impairment scale (AIS D, n=9; C, n=3; B, n=2; A, n=3) and initial neurological level of injury (C2 - T10, cervical=9, thoracic=8) (6 individuals did not complete the full study) (Fig. 7). Not all SCI subjects could be tested at each timepoint and at all intended spinal segments (at 1 month, n=14; at 3 months, n=11; at 12 months, n=8). The mechanical detection threshold (MDT) was not different between SCI subjects and healthy controls in L4 (right shin) and L5 (dorsal right foot) dermatomes as expressed by Z-scores <±2 throughout. However, incomplete SCI subjects displayed a lowered mechanical pain threshold (MPT) compared to healthy subjects at L4 and/or L5 dermatomes at 1 month up to one year post-injury (Fig. 7, Z-scores >2). At 12 months, of the 8 remaining incomplete individuals (AIS-B, C and D), 5 reported at- (n=1) or below-level (n=4) NP. In summary, incomplete SCI subjects are characterized by early and persistent hypersensitivity towards nociceptor mediated stimuli, whereas such changes could not be observed in respect to mechanoreceptor mediated stimuli.

**Figure 7:**
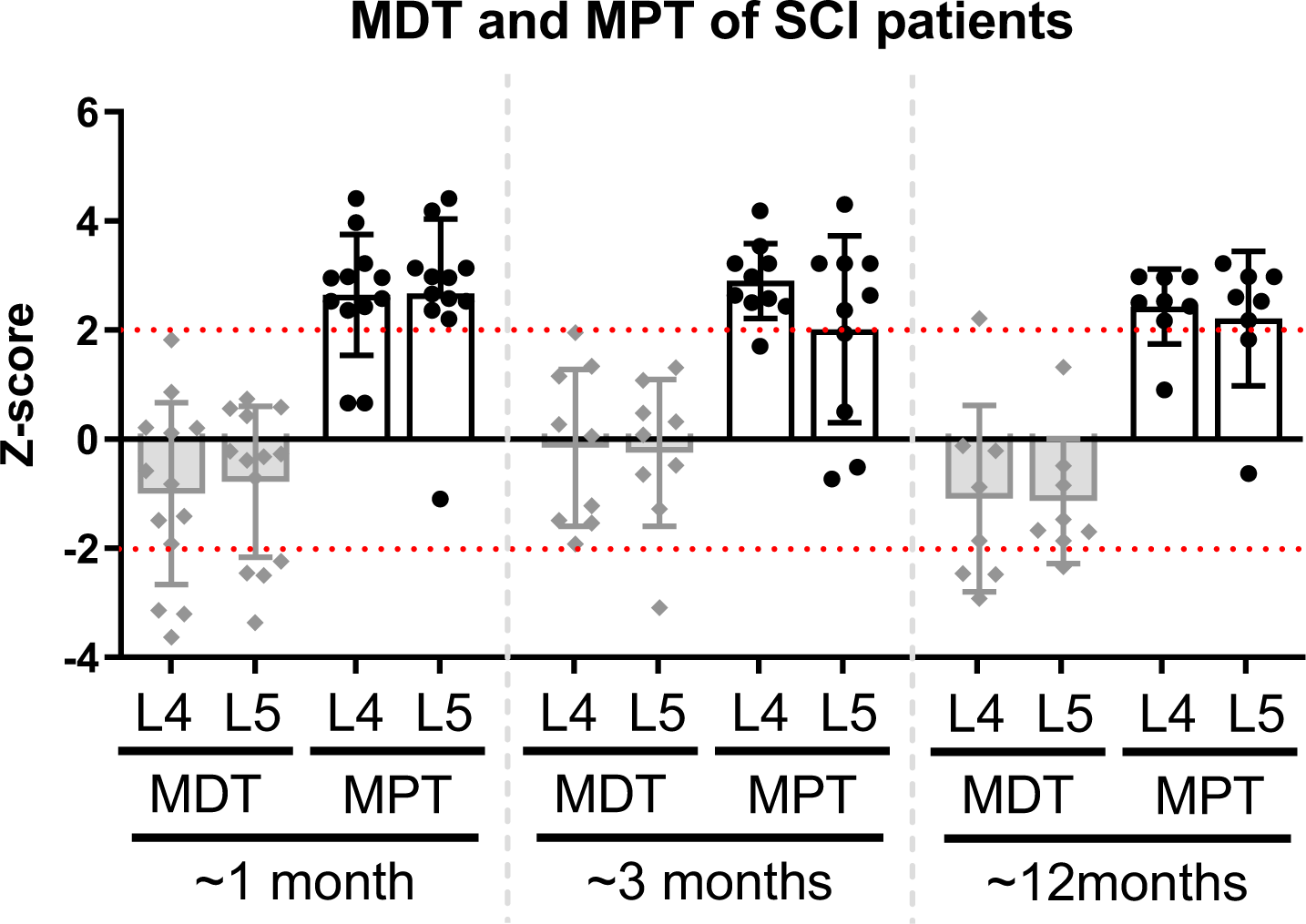
Quantitative sensory testing in SCI subjects. 14 incomplete SCI individuals were examined 1, 3 and 12 months post-SCI for mechanical detection threshold (MDT), reflecting Aβ mechanoreceptors, and mechanical pain threshold (MPT), reflecting Aδ nociceptors. SCI individuals were measured at the L4 (right shin) and L5 (dorsum of right foot) dermatomes and found to be altered from healthy controls in MPT but not MDT (represented by Z-scores). Red dashed lines visually mark the Z-scores at 2 and -2. Mean ± SD

## DISCUSSION

Results from the present study improve our understanding of initiation and maintenance of NP signals in traumatic SCI. NP ameliorating effects of mechanoreceptor ablation depend on the timing of ablation in relation to the time of injury. Only pre-SCI mechanoreceptor reduces pain behavior, suggesting a potentially NP inducing role. Replicating previously published evidence, nociceptors become hyperexcitable early after SCI. Respective signals are likely further augmented by early structural rearrangement of nociceptive nerve endings in the dorsal horn of the spinal cord below injury level, which help to maintain NP over time.

The finding that pre-SCI ablation of mechanoreceptors - not post-SCI ablation - reduces pain behavior, points toward a NP inducing rather than a NP maintaining effect. This is supported by the absence of mechanoreceptor hyperexcitability in hindpaw skin-nerve preparation 7 days after thoracic contusion SCI in mice. A recent publication investigating hyperexcitability in DRG in a skin-DRG-spinal cord preparation 1 day after mouse contusion injury made the identical observation of absent mechanoreceptor hyperexcitability (30). Central branches of mechanoreceptors form the lemniscal tract, which reach from their entry point into the spinal cord below injury level all the way to and across the spinal cord injury site. There is a vast body of evidence describing molecular changes in DRG neurons caused by lemniscal tract lesions (31, 32), which not only underlie the poor regenerative capacity of these axons, but may also be causally linked to nociceptor hyperexcitability. Macrophage migration inhibitory factor (MIF) released within DRG has been reported to signal nociceptor hyperexcitability after SCI (33). Whether this a consequence of lemniscal tract injury or related to systemic signals released after SCI irrespective of specific lemniscal tract injury has yet to be determined. A previous study, which would argue against the concept of retrograde signaling along the injured lemniscal tract to trigger NP behavior, reported reduced mechanical allodynia in a rat thoracic lateral hemisection SCI model after subsequent (1 day later) rostral dorsal column (containing the lemniscal tract/central branches of mechanosensory neurons) transection (34). Of course, animal species are different, a different SCI model and in particular a different approach to eradicate mechanosensory neuron derived signal transmission was chosen.

Post-SCI mechanoreceptor ablation indicates that specific input transmitted by mechanoreceptors, such as provided by spontaneous locomotion in the cage or augmented by treadmill training, has no effect on NP behavior in SCI mice. Not only are there no major changes in respect to pain behavior compared to non-ablated SCI mice, but enhanced peptidergic nociceptor sprouting in the dorsal horn post-SCI was also not altered by mechanoreceptor ablation. Nevertheless, we and others have identified a robust pain relieving effect of sensorimotor activity such as treadmill training (2, 3, 35, 36). Of course, other pain reducing mediators of activity/exercise have been identified, which do not rely on neural transmission through mechanoreceptors. For example, in a spinal nerve ligation model of NP, endogenous opioid content was increased in the brainstem following treadmill training, whereas such effects were reversed by opioid receptor antagonists (37). Following direct injury to the PNS - spared nerve injury (common peroneal and tibial nerve ligated and cut, sural nerve left intact) – identical transgenic mice, in which TrkB expressing mechanoreceptors were ablated - did not develop pain behavior (mechanical allodynia) (5). Of course, following experimental SCI, there is no direct mechanical injury to the peripheral branch of DRG neurons, which could account for the observed differences.

One week following moderate thoracic contusion SCI in mice, nociceptors were identified to be hyperexcitable in skin-nerve preparations (lower activation thresholds and afterdischarges). These findings are in line with studies in cultured DRG neurons (38–41) and confirm recent evidence in skin-nerve-DRG-spinal cord preparations taken 1 day post-injury in a moderate mouse contusion SCI model (30). Moreover, the observation of afterdischarges in nociceptors represents the likely correlate of prolonged hindpaw nocifensive behavior (licking, shaking, holding) in response to lighter von Frey hair filaments in SCI contused mice 7 days post-injury and goes beyond reflexive movements thus suggesting a supraspinal pain response.

Having observed aberrant sprouting of (peptidergic) nociceptors it was intended to apply an ablation technique to nociceptors to investigate a causal role in mechanical allodynia. In a PNS injury model (spared nerve injury), late ablation of nociceptors in SNS^iDTR^ mice fully abrogated mechanical allodynia of disorganized reinnervated tibial nerve regions (6). Unfortunately, the poor health condition of SNS^iDTR^ mice in our hands with the presentation of seizures was not sufficient to inflict contusion SCI and a prolonged survival period on these animals. In SNS^tdTomato^ we observed a leaky reporter expression (42) that based on breeding records could be traced back to Rosa26LSL-tdTomato mice. SNS^tdTomato^ animals exhibiting a wrong reporter expression presented with red paws and removed from the study. Even inner organs appeared red following transcardial perfusion. Of course, in SNS^iDTR^ mice the leaky expression would not be visible, because instead of the tdTomato reporter, the iDTR cassette was inserted into the Rosa26 locus and driven by the CAG promoter.

In subjects suffering from subacute traumatic SCI, hypersensitivity to mechanical pain stimuli was observed, which in principle is supported by preclinical findings of nociceptor hyperexcitability and dorsal horn nociceptor sprouting. Of course, in human subjects systematic assessment options are limited, which do not allow to clearly identify neuroanatomical regions (DRG, dorsal horn), where signals for NP are initiated and maintained. The observed hypersensitivity to mechanical pain stimuli was identified in SCI patients with and without NP. This points again towards co-factors such as the integrity of ascending sensory pathways within the spinal cord, which together with DRG hyperexcitability and dorsal horn nociceptor sprouting define NP presentation. Standardized sensory examination according to International Standards for Neurological Classification of SCI (ISNCSCI) in subacute traumatic and ischemic SCI patients observed within the European Multicenter Study about Spinal Cord Injury (EMSCI) revealed relative preservation of the spinothalamic tract (pinprick sensation), which predisposes for NP presentation (unpublished observation). In clinically sensorimotor complete SCI subjects (AIS-A), an instrumented analysis with contact-heat evoked measurements (CHEPS) also identified partial preservation of the spinothalamic tract to be associated with NP (43).

Taken together, mechanoreceptors are likely involved in the initiation, not the maintenance of NP. Nociceptors are confirmed as the main drivers of NP maintenance, which is highlighted by the hyperexcitability of nociceptors, nociceptor sprouting in the dorsal horn following experimental SCI complemented by nociceptor hypersensitivity in human SCI subjects. Hyperexcitability inducing factors, either transmitted systemically or retrogradely along injured axons (central branch of mechanoreceptors) into/towards DRG have yet to be defined. According to this understanding, the term “central NP” used to describe SCI-induced NP should be reconsidered. Of course, the initial trigger results from the central nervous system injury. However, the most likely driver of SCI-related NP sits in the peripheral nervous system showing also many similarities to pathophysiological changes observed following direct peripheral nerve injury.

## Supporting information

Sliwinski et al Suppl Figures

## ACKNOWLEDGMENTS

Authors would like to give thanks to Armin Blesch for personal communication and input. This research was funded by (1) Deutsche Forschungsgemeinschaft (SFB 1158) awarded to R. Puttagunta, N. Weidner, S. Lechner, P. Heppenstall and R.Kuner, (2) Olympia Morata Program Fellowship of the University of Heidelberg Faculty of Medicine awarded to R. Puttagunta, (3) Onassis Foundation scholarship for doctorate studies (ID:F ZM 034-1/2016-2017) awarded to V. Kampanis and (4) X. Cheng sponsored by China Scholarship Council (CSC).

## AUTHOR CONTRIBUTIONS

Conceptualization, R.P., N.W., C.S., and S.F; Methodology, C.S., S.G.L., P.A.H. and R.K.; Formal Analysis, C.S., R.P., L.H., and S.G.L.; Investigation, C.S., L.H., F.J.T., L.W., V.K., X.C., M.M., S.F, and S.G.L.; Writing – Original Draft, C.S., R.P., B.T., S.F. and N.W.; Writing – Review & Editing, C.S., R.P., S.G.L, N.W., S.F. and L.H.; Resources, N.W., R.K., and P.A.H.; Supervision, R.P., N.W., S.F., and S.G.L.; Project Administration, C.S., R.P., and S.F.; Funding Acquisition, N.W., and R.P.

## Supplementary Figure Legends

**Suppl. Fig. 1:**
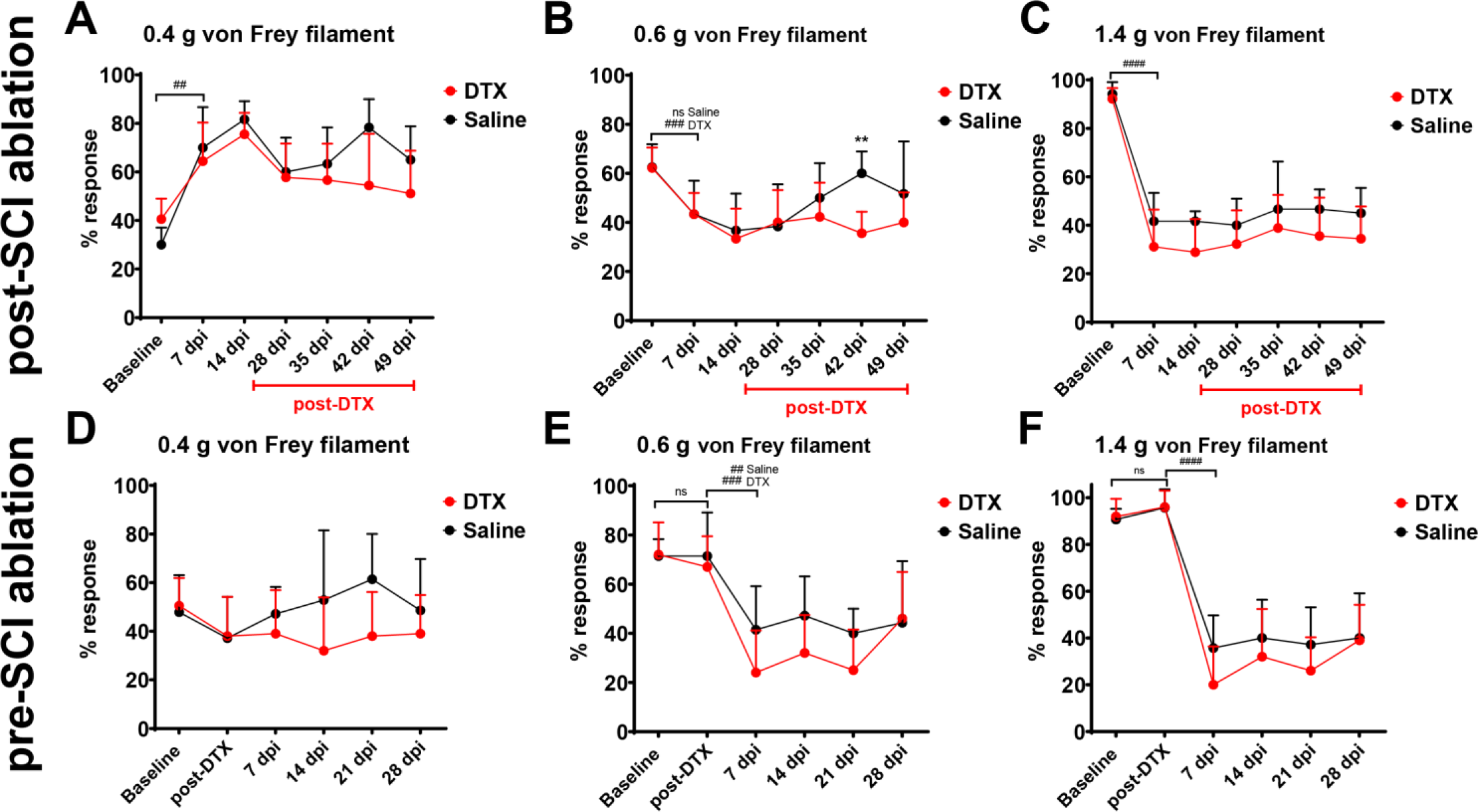
Mechanical sensitivity of 0.4, 0.6 and 1.4 g von Frey hair filaments following pre- and post-SCI TrkB-expressing mechanoreceptor ablation. For post-ablation of TrkB+ neurons, no overall group differences are observed for the 0.4 g filament (Two-way RM ANOVA group differences p=0.17, significant time difference between baseline and 7 dpi ## p<0.01) (**A**) and the significant injury-induced hyposensitivity (Two-way RM ANOVA time differences between baseline and 7 dpi ### p<0.001 and ns not significant) to larger-diameter noxious 0.6 g and 1.4 g filaments (**B, C**) remain relatively unaffected by ablation of TrkB+ sensory neurons (Two-way RM ANOVA, group differences 0.6 g, p=0.04; 1.4 g, p=0.01). Similarly, for pre-ablation of TrkB^+^ neurons, no overall group differences are observed for the 0.4 g filament (**D**) and the significant injury-induced hyposensitivity to larger-diameter noxious 0.6 g and 1.4 g filaments (**E, F**) remain unaffected by ablation of TrkB+ sensory neurons (Two-way RM ANOVA, group differences 0.4 g, p=0.11; 0.6 g, p=0.11; 1.4 g, p=0.13; time differences baseline vs post-DTX, ns not significant, post-DTX vs 7 dpi, ## p<0.01, ### p<0.001, #### p<0.0001). Mean ± SD

**Suppl. Fig. 2:**
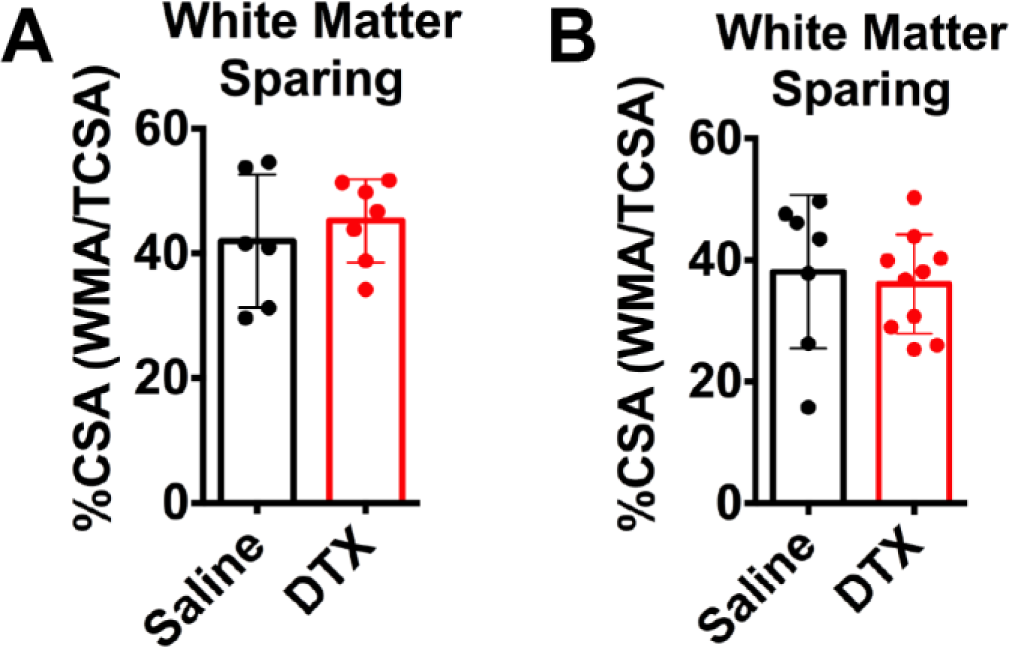
White matter sparing of T11 spinal cord sections following post and pre-ablation of TrkB-expressing sensory neurons. (**A**) Quantification of the cross-sectional area (CSA) at the lesion epicenter indicates the loss of gray and white matter. No difference in lesion size analyzed as white matter sparing is observed comparing saline-injected control with post-SCI ablated injured TrkB^DTR^ animals. (mean ± SD; Saline n=6; DTX n=7) (**B**) No lesion size difference is observed comparing saline-injected control with pre-SCI ablated injured TrkB^DTR^ animals. (mean ± SD; Saline n=7; DTX n=9)

**Suppl. Fig. 3:**
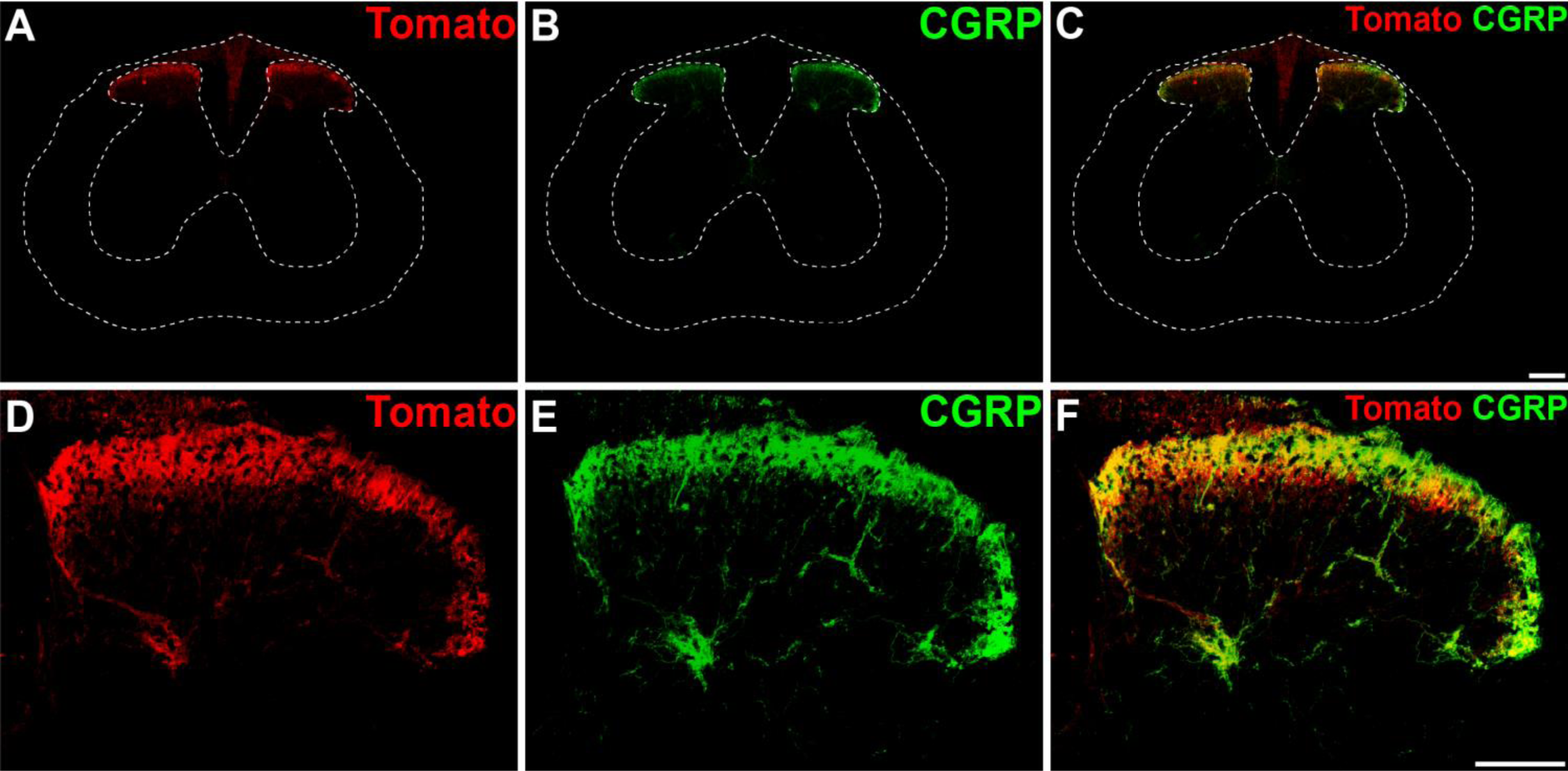
Termination pattern of tdTomato+ fibers in the lumbar spinal cord. Representative lumbar spinal cord sections showing tdTomato+ (**A, D**) and CGRP+ fibers (**B, E**) fibers in the lumbar (L4-L6) dorsal horn. In line with their function, they mostly terminate in the superficial laminae and largely overlap (**C, F**). (scale bar A-C, 200 μm; D-F, 100 μm).

**Suppl. Fig. 4:**
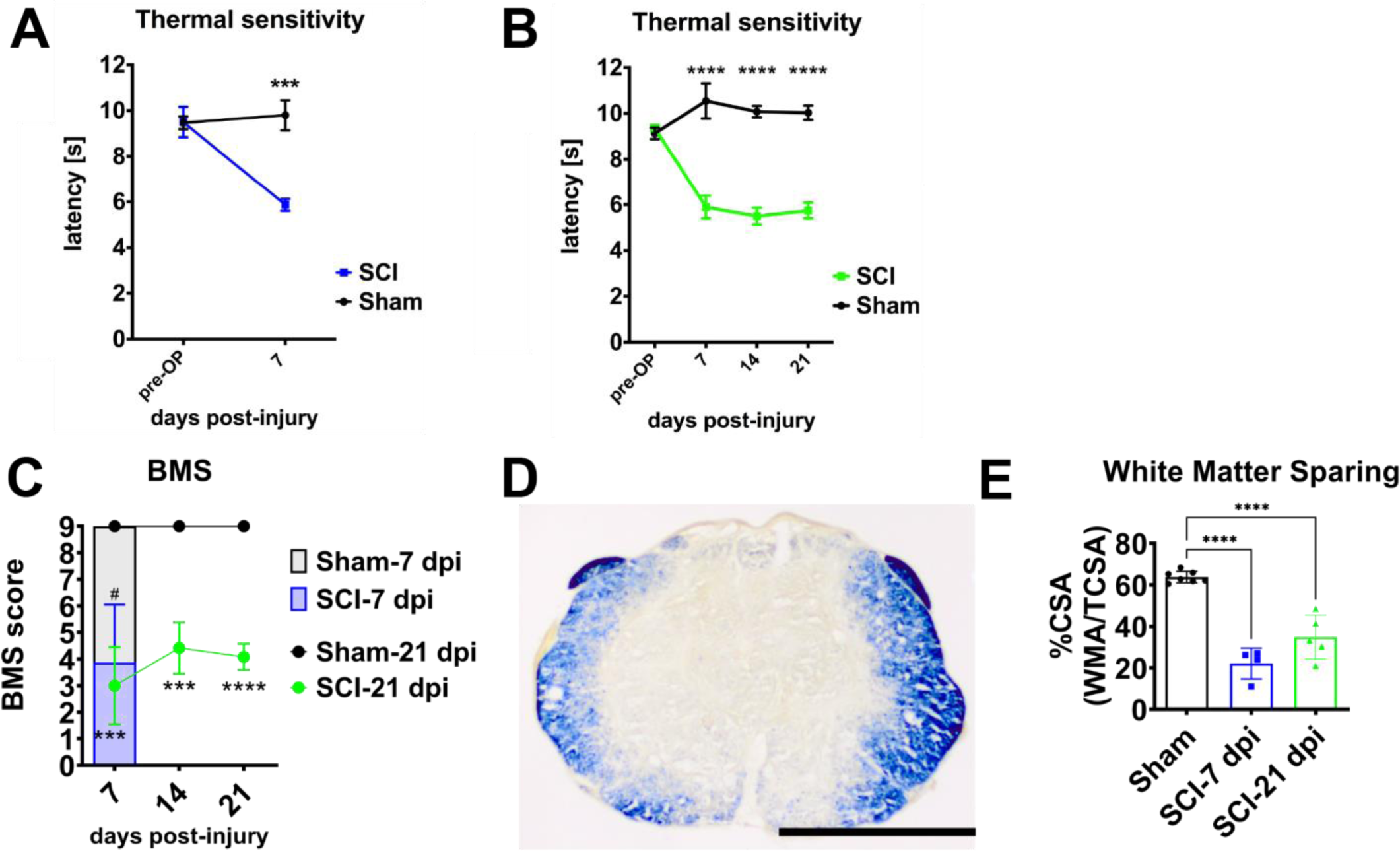
SCI-induced thermal hyperalgesia and motor deficits after white and gray matter damage in SNS^tdTomato^ mice. (**A**) SCI-7 dpi animals show a reduced response latency in the Hargreaves test. (**B**) A reduced response latency in SCI-21 dpi mice is already evident by 7 dpi and this thermal hypersensitivity is stable until the end of the experiment. (*** p<0.001, **** p<0.0001 comparing SCI with Sham animals) (**C**) T11 moderate contusion (50 kDyn) injury in SNS^tdTomato+^ mice leads to stable motor deficits over a 21 dpi period. 7 dpi group: Sham-7 dpi (n=3), SCI-7 dpi (n=4), unpaired t-test # p<0.05; 21 dpi group: Sham-21 dpi (n=4), SCI-21 dpi (n=6), Two-way RM ANOVA, Sidak pairwise comparisons, *** p<0.001, **** p<0.0001. (**D, E**) Lesion area analysis shows the stability of the lesion extent (One-way ANOVA compared to Sham, **** p<0.0001).

**Suppl. Fig. 5:**
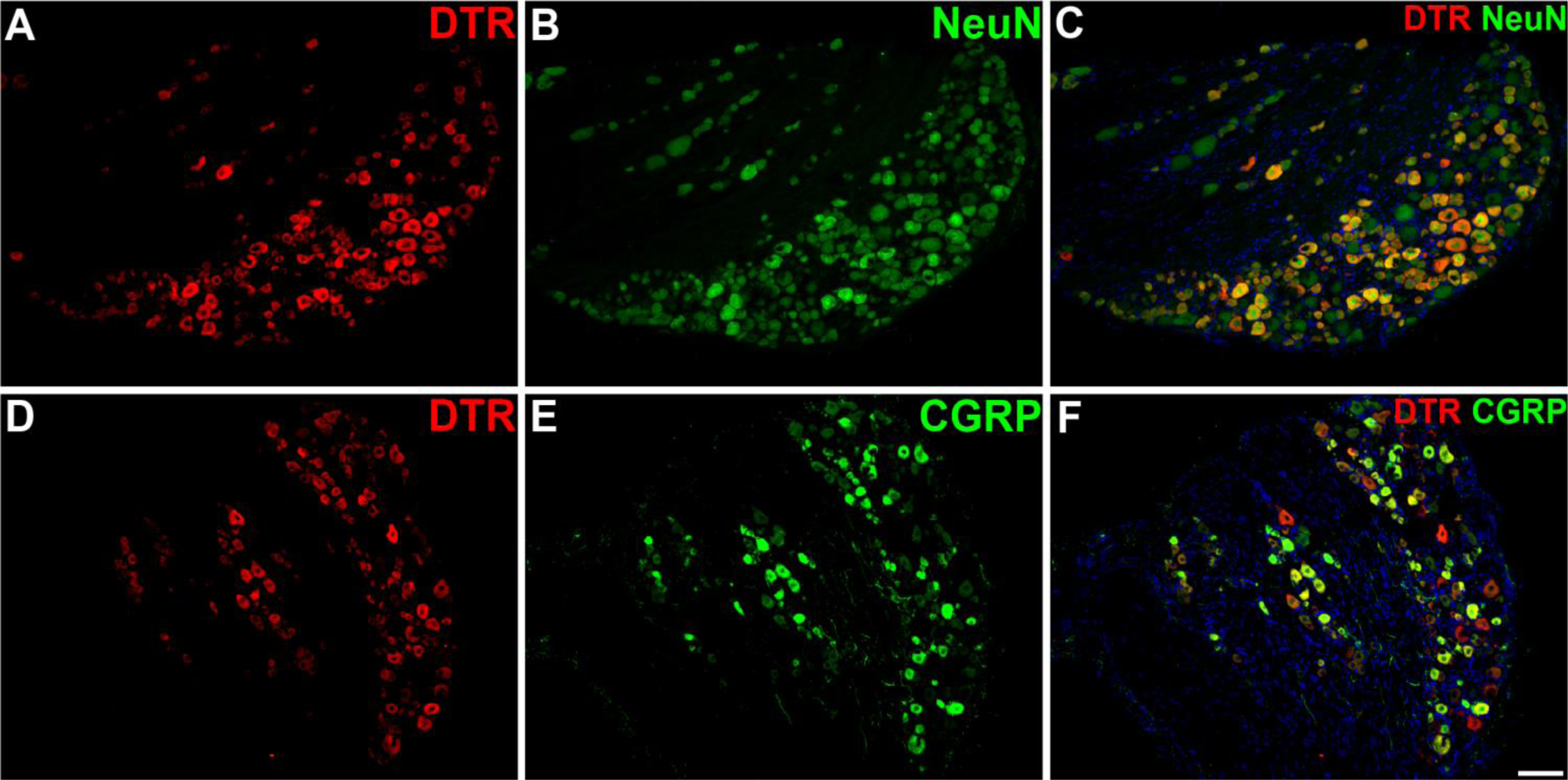
Lumbar DRG sections of SNS^iDTR^ mice for ablation of nociceptive neurons using diphtheria toxin. (**A-C**) Labeling of lumbar (L4-L6) DRG neurons of SNS^iDTR^ mice for human heparin binding EGF-like growth factor (HB-EGF), which functions as the diphtheria toxin receptor (DTR) and NeuN shows a similar expression pattern seen in SNS^tdTomato^ animals. (**D-F**) A large percentage of DTR+ sensory neurons co-express CGRP. (scale bar, 100 μm).

