## Supplementary figures and images for "Pre-Injury Mechanoreceptor Ablation Reduces Nociceptor-Driven Spinal Cord Injury-Induced Neuropathic Pain"

### Sliwinski et al Suppl Figures

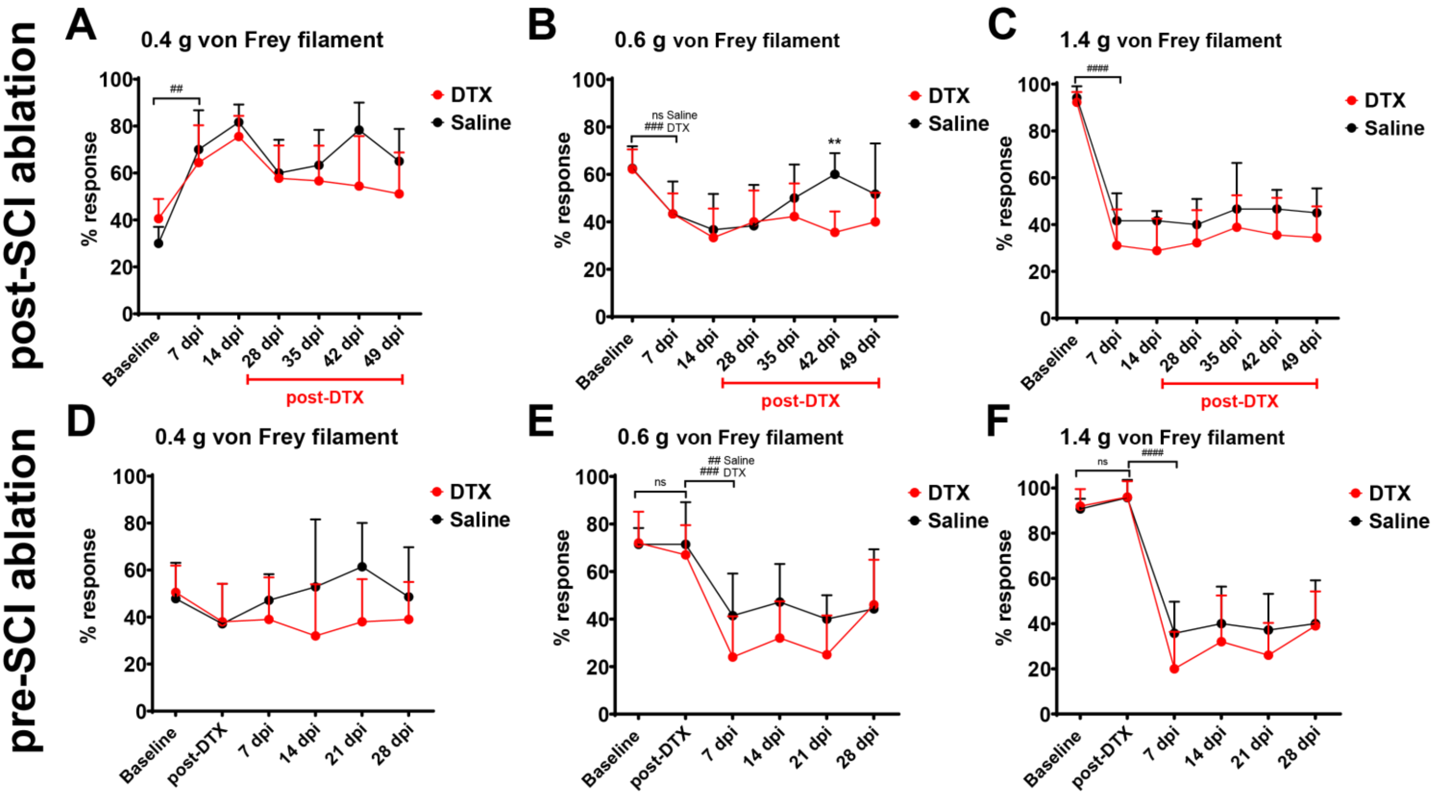

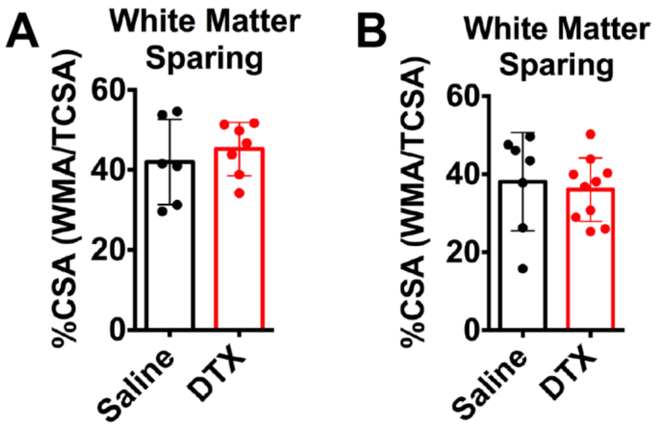

**Suppl. Fig. 3**

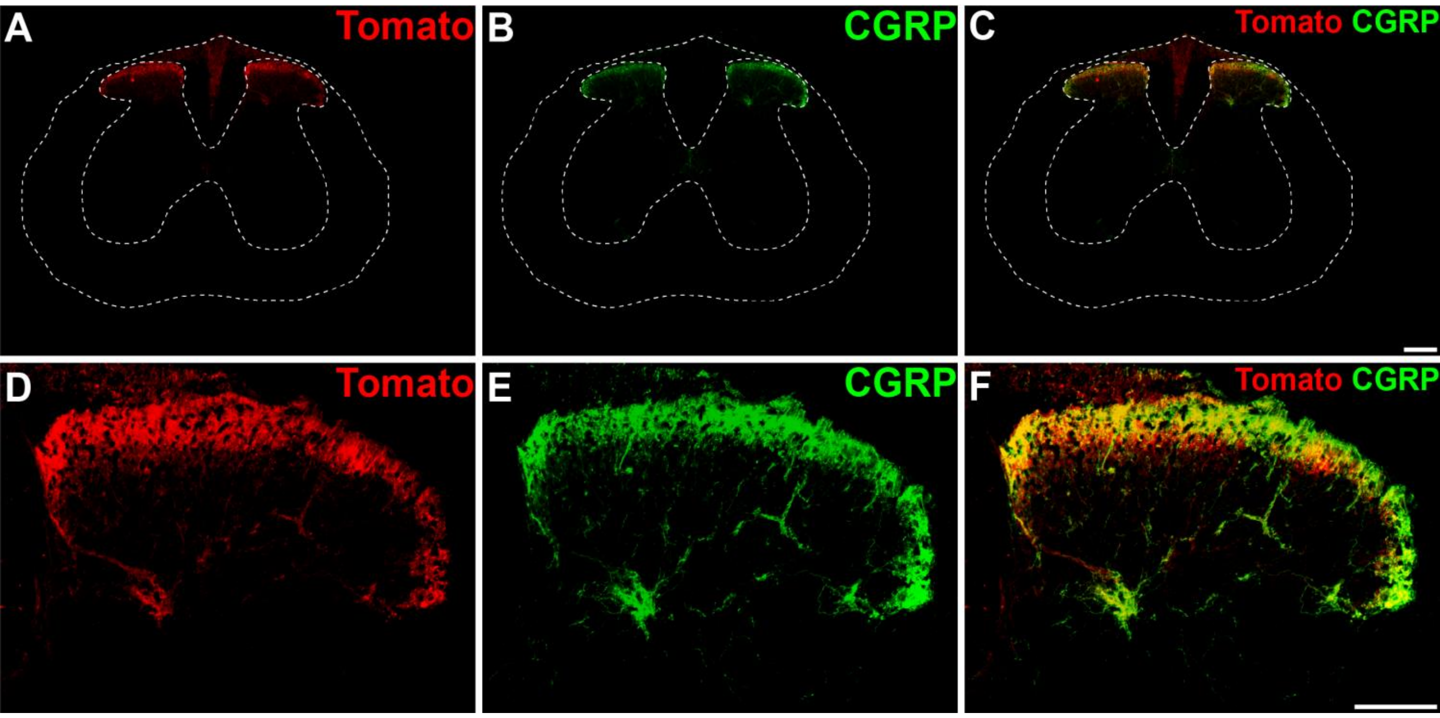

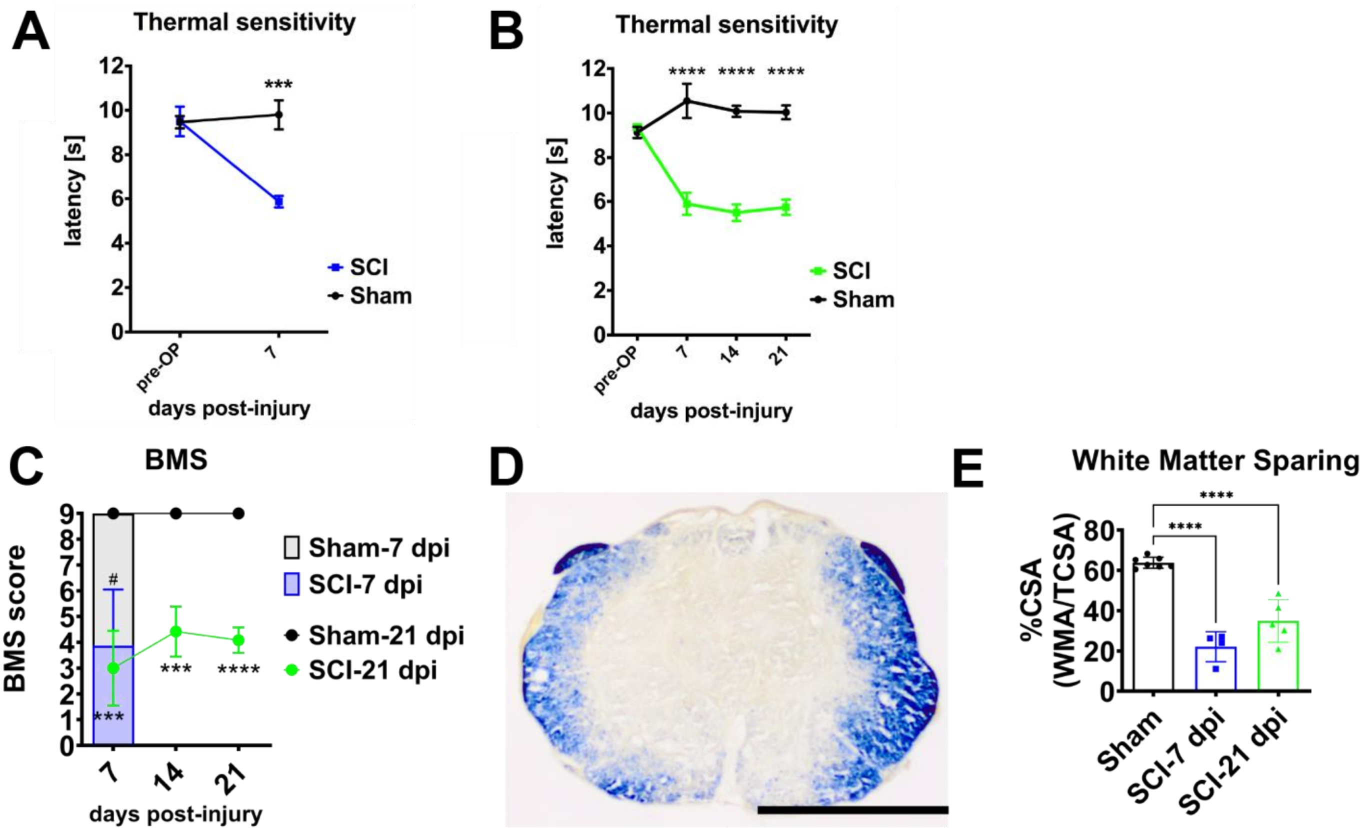

**Suppl. Fig. 5**

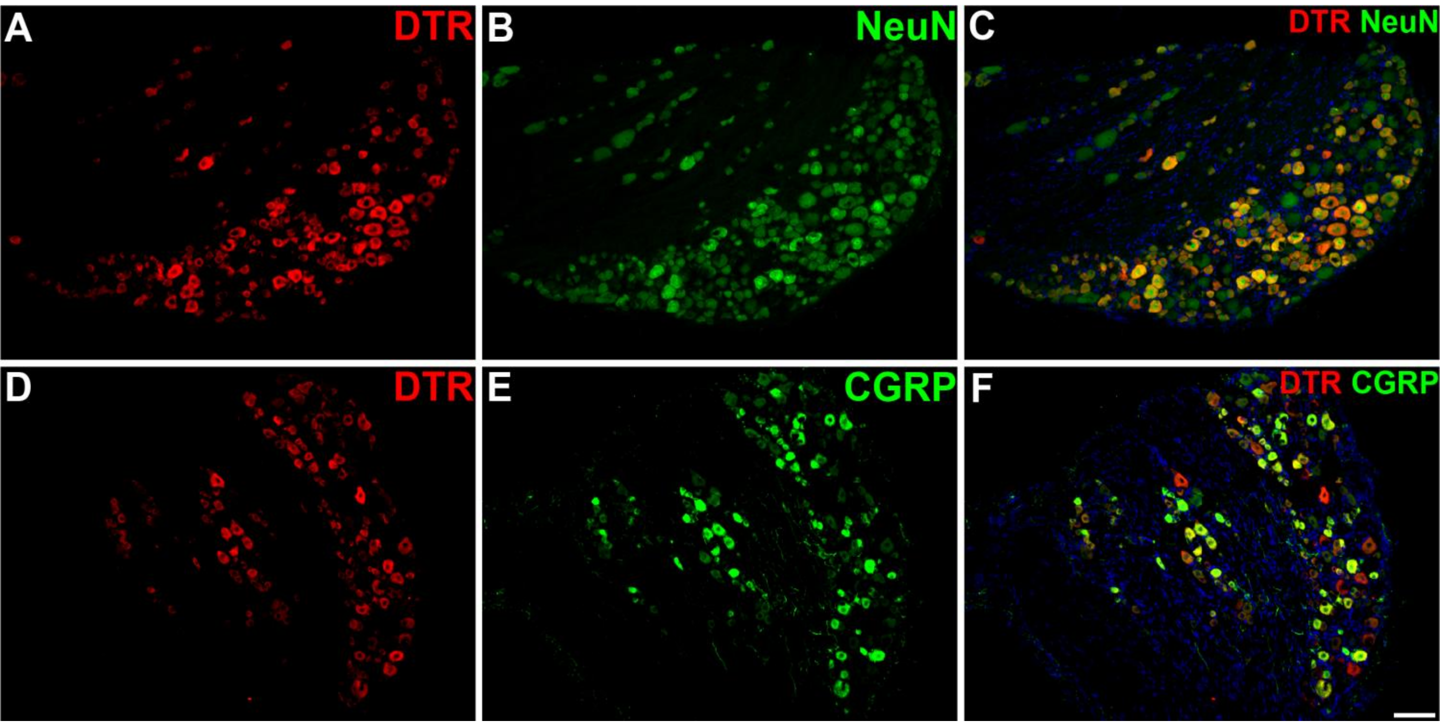
